# LYNX: a deep generative model for linking spatial dynamics and cell interactions in multimodal spatial data

**DOI:** 10.64898/2026.07.09.737574

**Authors:** Yinuo Jin, Joshua D. Myers, Presha Rajbhandari, Jia Yi Zhang, Kaylee W. Fang, John Shaw Moazami, Noreen Hosny, Brent R. Stockwell, Elham Azizi

## Abstract

Tissues are spatially organized systems in which cell states, functions and interactions vary across spatial coordinates, forming compartments or gradients shaped by local microenvironments. Understanding how molecular features and cell–cell interactions change across space and time is central to studying development, homeostasis and disease. Addressing these questions increasingly requires the integration of multi-modal spatial data, which provides complementary views of cellular and structural organization. However, existing computational approaches typically combine modalities by weighting them equally, overlooking domain-specific technical artifacts, differences in spatial resolution and non-overlapping feature spaces. In addition, methods for spatial cell–cell communication analysis are largely developed for single-modality settings and do not model how interactions vary across the tissue. To address these gaps, we introduce LYNX, a deep generative framework that learns a shared latent representation of spatial dynamics from joint-measured modalities in the 2D or 3D domain, to provide a unified coordinate system for modeling how cell–cell interactions, phenotypes, and molecular programs vary along continuous spatial gradients. LYNX identifies spatial programs difficult to resolve with existing approaches, including metabolically coupled porto-central interaction remodeling in liver, recovery of degraded proteomic signals along the cortico-medullary axis in thymus, and branching trajectories towards DCIS and invasive niches marked by distinct stromal activation-states and immune-tumor crosstalk in breast tumor microenvironment. We demonstrate that LYNX robustly infers spatially resolved gradients, maps functional compartments and cell-cell interactions along spatial axes and is compatible across diverse spatial profiling technologies, modalities, and resolution disparities. LYNX provides a foundational and scalable framework to advance our understanding of healthy tissue physiology and to decode temporal evolution of complex diseases.

## 1 Introduction

The spatial organization of tissues is a fundamental determinant of cellular behavior. Local microenvironmental cues including signaling molecules, nutrient availability, extracellular matrix composition, and physical constraints, shape cell states, functional programs, and patterns of cell–cell communication across tissue space. As a result, molecular activities and cellular interactions become organized into spatially structured programs that coordinate development, homeostasis, and disease [1]. In a structurally zonated organ like liver, blood flow along the porto-central axis establishes a graded microenvironment that directly governs the metabolic zonation and physiological functions [2–4]. Similarly, the intestinal epithelium exhibits a spatial gradient, wherein the epithelial cells undergo proliferation and differentiation along a continuous crypt-villus axis [5–7]. In both systems, key signaling pathways such as Wnt/*β*-catenin contribute to the spatial organization of cellular functions by regulating pericentral metabolic programs in the liver and stem-cell maintenance in the intestinal crypt. [8, 9]. Similarly, in the context of cancer, physical and chemical gradients, including oxygen levels, pH and chemokine signaling, profoundly influence heterogeneous tumor phenotypes with diverse migration and immune evasion patterns [10–12]. Together, these examples highlight how tissue architecture organizes molecular programs and cellular interactions across space. Characterizing these spatial patterns is therefore essential for understanding how microenvironmental context shapes tissue function in both healthy and diseased states. Moreover, linking molecular features to intercellular communication across tissue space is critical for understanding how local environments influence cellular behavior and collective tissue function.

The rapid advancement of spatial profiling technologies has created unprecedented opportunities to interrogate spatially organized biological systems at different resolutions. Diverse spatial modalities such as probe- or sequencing-based spatial transcriptomics [13, 14], mass spectrometry imaging based spatial metabolomics [15, 16], highly-multiplexed antibody based imaging based targeted proteomics [17–20], and spatial epigenomics [21] capture complementary molecular maps of tissue organization. By preserving the spatial organization that dissociative single-cell workflows typically lose, these technologies are now capable of providing rich information on cellular organization and molecular distribution in healthy and complex diseased microenvironments. Data obtained from multimodal spatial measurements from the same or adjacent tissue sections now offer opportunities to obtain holistic views of tissue architecture that links transcriptomic, proteomic, metabolic, epigenomic, and histological information within a shared spatial framework [22].

Computational approaches to gain biological insights on spatio-temporal dynamics from these complex and high dimensional datasets have lagged behind the rapid advancements in spatial technological developments. The challenges are three-fold. First, many spatial analysis methods focus primarily on identifying discrete spatial domains or regions within tissues [23–25]. While such approaches are well-suited for delineating boundaries, they fail to model tissue organization exhibiting continuous spatial variation, gradual state transitions, and recurrent expression programs that do not conform to rigid cluster assignments. This limitation is especially important in the context of development, tissue morphogenesis and disease progression [10, 26]. Second, most approaches operate on a single modality, as integrating multiple modalities to recover coherent, continuous spatial organization remains computationally challenging. It stems from differences in cross-scale alignment of images acquired at different spatial resolution, number of features, and technology specific artifacts. These mismatches complicate the construction of a shared representation that preserves biologically meaningful spatial gradients and enables downstream analysis of gradient-resolved molecular programs and cell–cell interactions. Furthermore, current vertical integration approaches such as multi-view autoencoders, Product-of-Experts (PoE), and feature concatenation [27–29] can obscure modality-specific signal or fail to distinguish technical artifacts (e.g. transcriptomic dropout, autofluorescence, noise and degradation in proteomic imaging [30], and hotspots in metabolomics imaging [31]) from biological signal shared across modalities. More importantly, a growing class of methods have been proposed to infer cell–cell communication from spatial data ([32–35]), typically leveraging spatial proximity and ligand–receptor structure. However, these approaches are often developed in single-modality transcriptomic settings and, critically, do not couple communication inference to spatial gradients and tissue architecture. This limits the ability to quantify how cellular interactions reorganize across microenvironments varying in metabolites, nutrients, cytokines, or other cues. Together, these limitations highlight the need for a unified framework that integrates heterogeneous multimodal spatial measurements, learns continuous spatial organization without predefined tissue compartments, and quantifies how cellular phenotypes and cell–cell interactions shift along the spatial gradients.

To address these challenges, we introduce LYNX, a hierarchical deep generative model for uncovering tissue gradients linked to spatially dynamic molecular programs and rewiring of cellular interactions, through integration of multimodal spatial expressions. LYNX is explicitly designed to accommodate heterogeneous spatial resolutions, non-overlapping feature spaces, and non-homogeneous assay-specific technical noise without predefined tissue axes. By employing cross-attention mechanisms on a joint-modality heterogeneous graph to map paired spatial measurements into a shared embedding space, LYNX enables the combined analysis of spatial dynamics in molecular features and variation in cell–cell interactions across spatial contexts. As a scalable representation learning approach for high-resolution spatial datasets, LYNX connects complementary features spanning transcriptomes, proteins, lipids, and metabolites within the spatial coordinate to enable the identification of spatially coordinated molecular programs and interaction networks, shaping tissue organization across diverse biological systems and disease states.

## 2 Results

### Overview of LYNX

LYNX is a deep generative framework that learns a shared latent representation from multimodal spatial data to enable the inference of continuous spatial gradients together with associated molecular and cellular dynamics. By incorporating data-driven priors and a graph attention mechanism within a hierarchical generative architecture, LYNX performs multimodal alignment and distance-aware modeling of cell-cell interactions in a unified probabilistic framework. This design facilitates downstream characterization of spatial organization across both niche and cellular resolution.

Given spatial modalities from adjacent tissue sections, LYNX takes a primary observation *x*^*N*×*G*^ and an auxiliary observation *u*^*M*×*P*^ as inputs. By convention, the primary observation corresponds to a higher-resolution or higher-throughput modality, e.g. spatial transcriptomics measuring gene expression in *N* cells (or spots), whereas the auxiliary modality may have coarser spatial resolution (*M < N*; e.g. patch-level spatial metabolomics imaging), or fewer features (*P* ≪ *G*; e.g. spatial proteomics or histology images). This formulation accommodates a wide range of heterogeneous modality pairings, e.g. high-resolution spatial transcriptomics with lower-resolution metabolomic imaging, transcriptomic–proteomic measurements with differing feature dimensionality, and transcriptomic–histology integration (**Fig. 1a**). As a preprocessing step, a heterogeneous spatial graph is constructed to link neighboring readouts within and across modalities upon image registration (**Fig. 1b, Fig. S1a**, see **Methods**).

**Figure 1:**
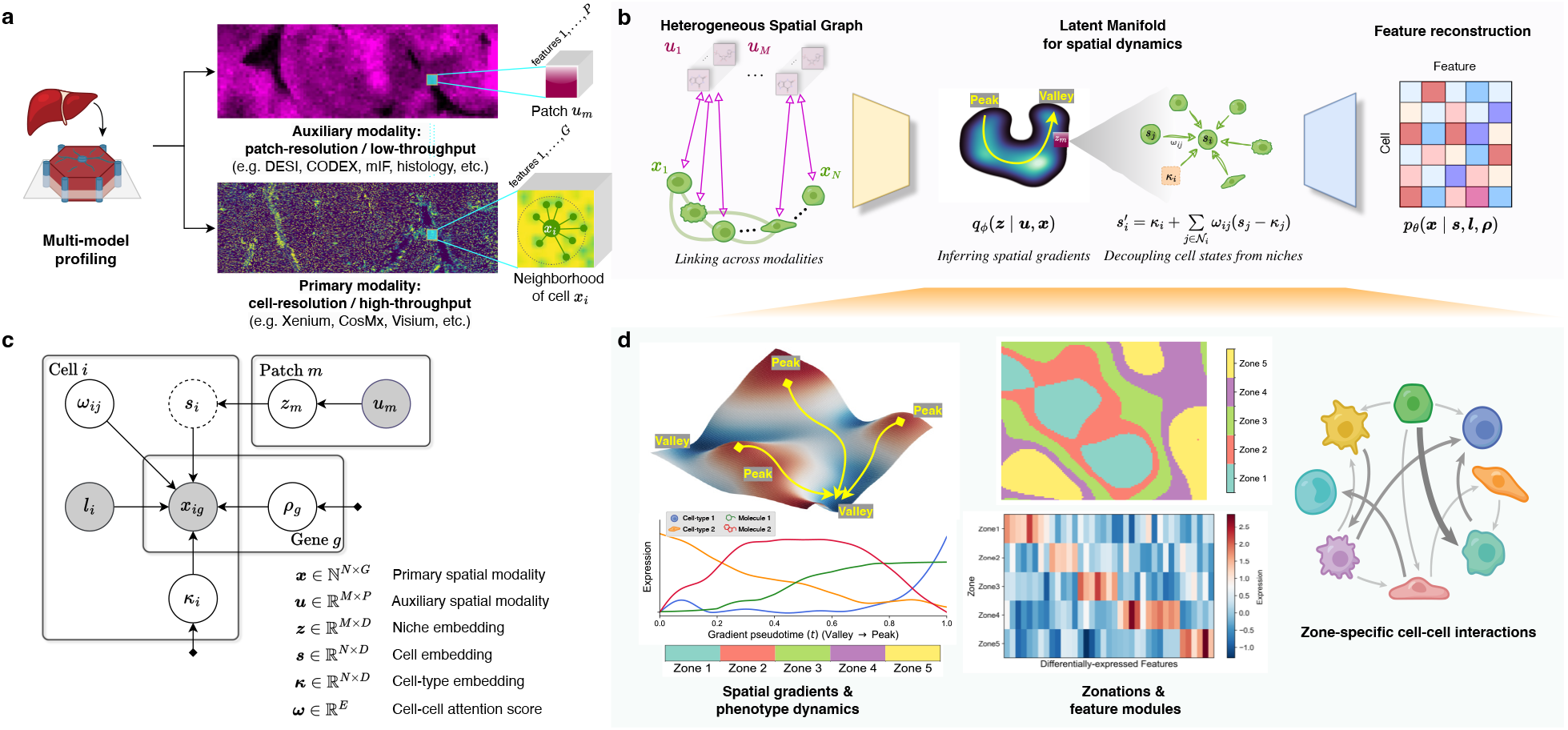
The LYNX model overview. **a**, LYNX takes paired multi-omics spatial data from adjacent tissue slides as input, allowing modalities to differ in resolution. **b**, Schematic representation of the LYNX architecture. From left to right: (i). LYNX employs a heterogeneous graph to connect spatial neighbors within and across modalities. (ii). A Graph Attention Network (GAT) encoder *q*_*ϕ*_ projects the paired expressions into a joint patch-level niche embedding (z). (iii). Unpooling and sparse message passing decouple cell-level embedding (s) from the effect of neighboring cells (**Methods**). (iv). An MLP decoder *p*_*θ*_ reconstructs the primary modality expressions (x) from the cell embedding. **c**, Plate model of the LYNX generative process; white nodes: latent random variables; shaded nodes: observation and auxiliary data; dashed nodes: deterministic variables. **d**, Downstream analysis. **Left**: spatial gradients (continuous) and zonation (discrete) predictions followed with corresponding phenotype dynamics and differentially expressed features along the inferred gradient coordinate; **Right**: cell-cell interactions across zones.

Utilizing a hierarchical variational autoencoder framework [28, 36–38], LYNX learns latent representations that capture shared spatial structure while preserving modality-specific variation (**Fig. 1b**). To infer the joint latent representation *z*, paired observations are projected into a niche-level embedding via cross-modal graph attention (**Fig. S1b,c**, see **Methods**). A niche refers to the local region surrounding a cell defined by a cell-cell graph. The niche embedding represents a shared microenvironmental context. In the generative process, a refined cell-level embedding *s* is obtained by deterministic unpooling of *z* to cell resolution, producing cell-level representations anchored in local spatial context. Unlike *z, s* is not sampled as a random variable; instead, it is computed through a deterministic mapping that propagates niche-level information back to individual cells. Together, the niche- and cell-level embeddings {*z, s*} define the latent manifold supporting downstream analyses.

A key distinguishing feature of LYNX compared to other multi-modal integration approaches is its conditional generative process in the decoder. Inspired by interpretable latent variable models [39–41], we utilize a coarse-to-fine generative design that places the lower-resolution/throughput auxiliary readout to shape a conditional prior over niche-level latent embeddings. Then, it deterministically refines to a cell-level embedding that generates the primary modality. The coarser auxiliary information of tissue architecture thus guides the latent structure without explicitly being reconstructed, while the model concentrates the generative capacity on the fine-grained primary readout. The process avoids noise propagation and mismatched likelihood contributions across modalities. The full hierarchical generative path is summarized in **Fig. 1c** (see **Methods: Generative process**).

Another distinguishing feature of LYNX is its explicit generative formulation that models both intrinsic cell states and spatially structured interaction effects. Specifically, we design a distance-aware sparse message passing mechanism to model cell-cell communication within the primary modality; similar to our previous work AMICI [35]. Following the aforementioned deterministic unpooling that obtains the cell-level latent *s*, LYNX further decomposes cell states into a baseline cell-type embedding *κ* and an interaction-associated residual derived from neighboring cells within the niche, such that 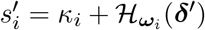, where *δ*^′^ denotes the mean-centered residual field *s* − *κ* (**Fig. 1b**, see **Methods**). The baseline term *κ* can be interpreted as intrinsic cell-type identity in the absence of cell-cell interactions, whereas the second term (*H* (·)) captures residual niche-associated deviations by applying a low-pass graph filter to neighboring residuals ((*s*_*j*_ − *κ*_*j*_)) using weights *ω*, thus representing spatially structured cell-cell communication (**Fig. S1e**, see **Methods**) [35, 42]. The residual weights *ω* are sampled from distance-informed priors, reflecting an underlying assumption that cells closer to each other are more likely to interact than their farther counterparts [35] (see **Methods**). Theoretically, increasing the distance penalty produces progressively sparser interaction neighborhoods, concentrating interaction weights on the nearest spatial neighbors (see **Methods**). This residual interaction module therefore links the spatial communication model to the broader LYNX generative framework: *u* conditions the niche-level latent representation *z*, which is deterministically unpooled to the cell-level latent *s*, adjusted through inferred intrinsic embeddings *κ* and residual weights *ω* to produce the interaction-adjusted state *s*^′^, and decoded to reconstruct the primary modality *x*. The corresponding variational inference path approximates the posterior over these latent components through cross-attention-based multimodal integration and residual interaction inference (**Fig. S1d**).

The core feature of LYNX is the interpretation of the learned embeddings for diverse downstream applications (**Fig. 1d**). First, spatially-informed gradients are derived from niche-aware latent embeddings via principal graph inference [43–45]. Subsequently, depending on the input data types, the end-to-end analysis can reveal the spatial dynamics of gene expression, or abundance of cell types, proteins, metabolites, etc., linked to the spatial gradient, highlighting spatially varying cellular phenotypes and molecular features. LYNX can also partition the continuous gradients into discrete and ordered functional zones, where zone-specific feature modules are subsequently identified (see **Methods**). Additionally, the learned weights (*ω*) provide quantitative summaries of cell-cell communication strength, and how they vary across spatially-organized zones or continuous gradients.

### LYNX uncovers metabolic gradients and zonation-specific cellular interactions in human liver

To evaluate whether multimodal spatial integration can uncover molecular and cellular programs associated with tissue organization, we first applied LYNX to human liver, a canonical example of spatial zonation organized along the portal vein (PV) to central vein (CV) axis, driven by graded portal-derived oxygen, nutrient and morphogens driving metabolic and functional zonation [2, 4, 8]. We profiled adjacent tissue sections from a healthy human liver with single-cell spatial transcriptomics using the Xenium platform (resolution: ~0.25 µm) [19] and meso-scale spatial metabolomics using desorption electrospray ionization (DESI) mass spectrometry imaging (MSI) (resolution: 40 µm) [46]. After preprocessing and alignment, 60,562 cells across 377 genes and 9,564 metabolic pixels spanning 615 mass-to-charge (m/z) channels were obtained from Xenium and DESI-MSI, respectively (see **Methods**). The 160-fold resolution disparity and non-overlapping feature spaces posed substantial challenges for multimodal integration methods, particularly those that typically require matched spatial resolution or shared features.

To establish a ground truth spatial axis, we performed multiplexed immunofluorescence imaging with metabolic zonation-specific antibodies on the same tissue section post Xenium and DESI image acquisition, used as held-out validation (**Fig. 2d**). To construct continuous labels, we derived a physics-informed gradient by solving a reaction-diffusion equation over a Delaunay triangulation spatial graph, with vascular landmarks from the antibody images as boundary conditions [47, 48] (see **Methods**). Since real lobular boundaries have irregular morphologies with unevenly distributed vascular positions across the liver parenchyma, this formulation captures non-linear spatial variation more accurately than distance-based heuristics, providing a quantitative reference for benchmarking.

**Figure 2:**
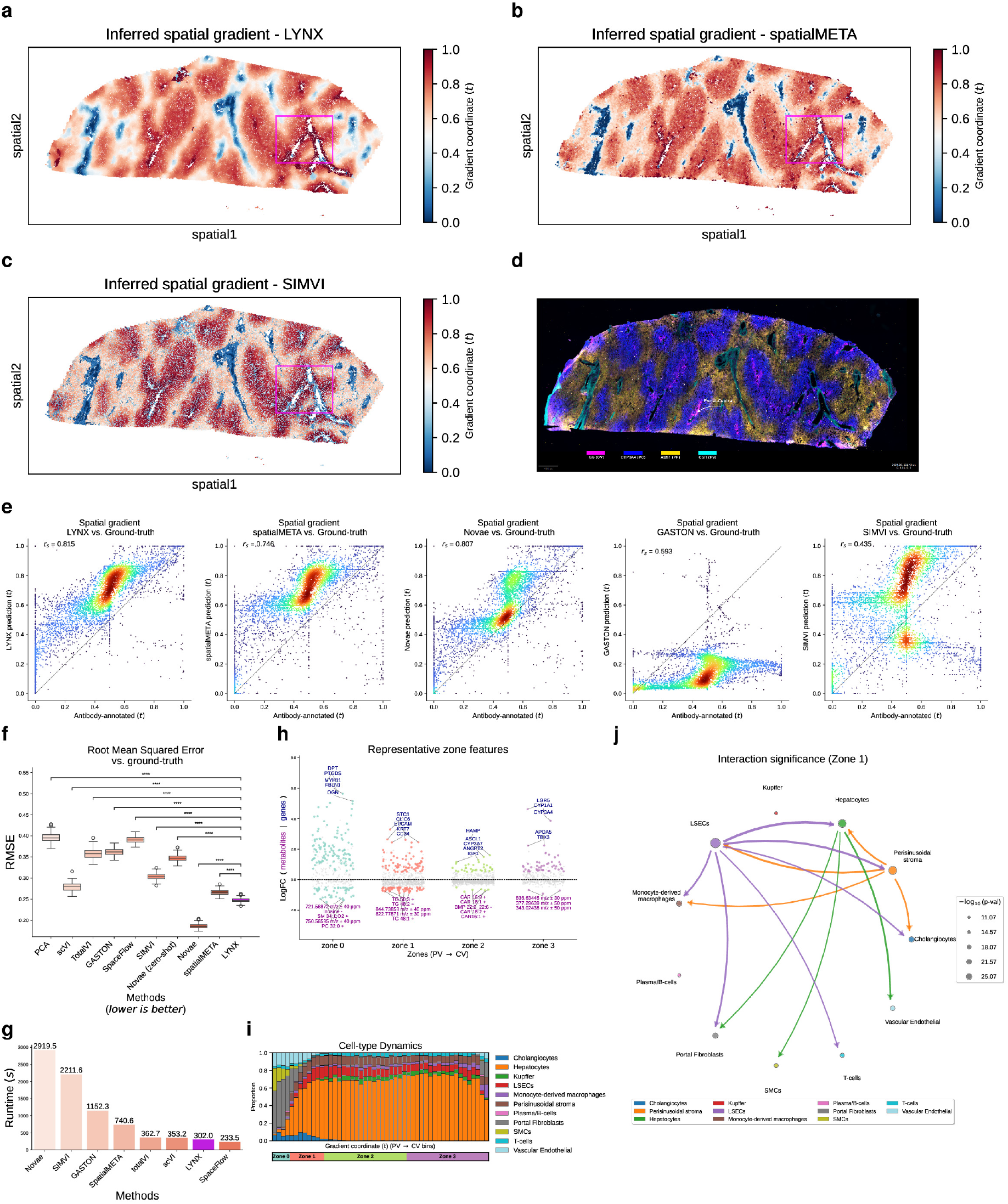
LYNX reconstructs spatial gradients and the underlying phenotype interaction dynamics along the porto-central axis in the human liver. **a-c**, Spatial gradients inferred along the portal-central axis by LYNX **(a)**, SpatialMETA **(b)**, and SIMVI **(c)**. The gradient coordinate (*t*) is scaled from 0 (PV end) to 1 (CV end) to enable direct comparison. PV: portal vein; CV: central vein. **d**, Experimental validation via antibody staining using zonation-specific markers. One representative porto-central axis between adjacent PV and CV is highlighted. **e-g**, Quantitative evaluation of the spatial gradients inferred by LYNX and alternative methods against the proxy ground truth derived from antibody staining (**Methods**): Spearman Correlation Coefficient **(e)**; root-mean-squared error (RMSE) **(f)**; and inference runtime across methods **(g)**. Statistical significance is computed with a one-sided independent t-test, **** *p <*1e-4; boxplots indicate the median (center lines), interquartile range (hinges) and 1.5× interquartile range (whiskers). **h**, Summary of representative upregulated genes and metabolites along the LYNX predicted porto-central axis. Zones are defined by discretized LYNX gradient coordinate, with zone 0 nearest the PV end and zone 3 nearest the CV end (**Methods**). **i**, Summary of phenotype dynamics predicted porto-central axis smoothed into equidistant bins. **j**, Cell–cell-interaction summary in zone 1 (periportal); Edge colors correspond to sender nodes, and widths represent interaction significance (**Methods**).

We benchmarked LYNX against the state-of-the-art methods for spatial gradient inference, single-cell and spatially-aware representation learning, and multimodal integration [36, 37, 49–53]. Spatial gradient visualizations reveal that LYNX produces gradients that are more consistent with antibody staining than competing methods (**Fig. 2a–d, Fig. S2a**). Quantitatively, LYNX achieved higher Spearman’s correlation (*r*_*s*_) and lower root mean squared error (RMSE) relative to ground-truth (**Fig. 2e,f, Fig. S2b**). Average precision (AP) and Receiver Operating Characteristic (ROC) were calculated against individual antibody intensities as well (**Fig. S2c,d**; see **Methods**). Across all metrics, LYNX significantly outperformed competing methods (**Fig. 2f**; one-sided *t*-test, *p <* 10^−4^), while maintaining superior runtime efficiency (**Fig. 2g**). In addition, our usage of graph layers led to spatially coherent gradients without fragmentation or over-smoothing (**Fig. S2a,e**). Notably, a key advantage of LYNX is its native ability to model modalities at drastically different spatial resolutions, whereas integration methods enforcing matched resolution (e.g., SpatialGLUE) failed to run. A similarly performing method to LYNX is Novae [52], a spatial foundation model pretrained on extraordinarily large-scale spatial transcriptomics datasets (CosMx, MERSCOPE, Xenium) across 18 human organ tissues, including liver, with substantially larger cell numbers and gene throughput. However, its application required large-scale pretraining on similar tissue domains and disease states, and incurs a runtime of approximately 10-fold higher than LYNX at the inference stage alone (**Fig. 2g**). Additionally, without fine-tuning on our data, Novae’s performance via zero-shot transfer learning was substantially degraded across all metrics (**Fig. S2a,d**). Beyond recovering continuous gradient, Novae’s built-in domain assignments produced spatially fragmented zonation, particularly between metabolic zones 2 and 3, whereas LYNX recovered coherent, contiguous lobular zones (**Fig. S3a**). Finally, as standard diagnostics for generative models [36, 37], we also verified that LYNX faithfully captures the primary modality via expression reconstruction accuracy (Pearson correlation: 0.907) with a meaningful latent manifold encoding the porto-central gradient (**Fig. S3c,d**).

Having established accuracy advantages across all benchmark metrics, we next leveraged the inferred gradient as a unified spatial reference to characterize how molecular programs and cellular interactions are organized across the hepatic lobule, enabled by curated marker-based cell-type annotations (see **Methods**). At the cell-population level, the inferred gradient revealed coordinated spatial organization of both parenchymal and non-parenchymal cell populations across the lobule. Cell-type abundance profiles recapitulated known architectural features of liver zonation while providing a unified spatial coordinate for examining multicellular organization(**Fig. 2i**; **Fig. S3e**). Cholangiocytes and smooth muscle cells (SMCs) were sharply concentrated in periportal regions, reflecting their anatomical confinement to the portal triad [54]. Notably, liver sinusoidal endothelial cells (LSECs) exhibited broad spatial organization spanning periportal and pericentral regions, positioning them as a potential intermediary between sinusoidal and hepatic counterparts, across the gradient. In addition, liver-resident kupffer cells which are reported to enrich periportally in mouse [55, 56], showed increasing abundance shifted towards midzone regions in our human data, consistent with a previous Visium study of human liver (**Fig. 2h, Fig. S3f**) [54], potentially reflecting complex heterogeneity in hepatic macrophage organization.

Within the macrophage community, reciprocal trends of enrichment between *CD68* and *MARCO* indicate polarized distributions of *MARCO+* and *MARCO-*macrophages from peri-portal to peri-central regions (**Fig. S3f**), suggesting a graded anti-inflammatory to pro-inflammatory environment along the porto-central axis [57]. T cells and plasma/B cells exhibit sparse and relatively stable distributions across zones consistent with the homeostatic immune surveillance in healthy tissues.

We next partitioned the continuous PV-CV gradient into four ordered zones following the hepatic zonation convention: zone 0 captures the portal triad-proximal region immediately surrounding the portal vein, while zones 1–3 correspond to the three canonical metabolic zones [8] spanning periportal (zone 1), mid-lobular (zone 2), and pericentral (zone 3) territories (**Fig. S3a**) and identified zone-specific molecular programs spanning both transcriptomic and metabolomic features (**Fig. 2h**, see **Methods**). Zone 0 was marked by stromal and smooth muscle markers *MYH11, DES*, and *CNN1*, consistent with portal triad SMC and fibroblast enrichment [54], alongside the purine metabolite inosine, experimentally validated as periportal-enriched in human liver [7]. The mid-lobular zone 2 was distinguished by *HAMP* (hepcidin), consistent with its established non-monotonic expression peaking at this zone [56]. Pericentral zone 3 was marked by well-known drug and xenobiotic metabolism markers *CYP3A4* and *CYP1A1* [2, 8] and phosphatidylcholine lipids (*PC 36:1*) validated with mass-spectrometry imaging [58]. Notably, *LGR5* emerged as one of the most enriched genes in the pericentral region, consistent with its hypoxia-induced pericentral expression. *LGR5* encodes a Wnt co-receptor that amplifies Wnt/*β*-catenin signaling, stabilizing *β*-catenin and driving peri-central transcriptional programs critical for liver regeneration, glutamine synthesis and other essential transcriptional programs in pericentral regions [8, 56, 59].

Having established coordinated molecular zonation, we next asked whether cellular communication networks are similarly organized across the porto-central axis. LYNX revealed substantial remodeling of intercellular communication along the lobular axis, indicating that liver zonation is accompanied by continuous changes in multicellular interaction networks rather than solely changes in molecular abundance. (**Fig. 2j**; **Fig. S3b**). Rich interaction diversity from the periportal region gradually diminishes toward the peri-central directions, consistent with graded sinusoidal supply of oxygen, nutrients, and portal-derived morphogens [8]. Lining the hepatic sinusoids and serving as the primary paracrine interface with hepatocytes [3, 60], LSECs emerged as the dominant sender in zone 1 actively communicating with a variety of hepatocytes and non-parenchymal cells (T-cells, fibroblasts and macrophages), suggesting roles in zone-specific hepatocyte imprinting and hepatic immune modulation consistent with their reported angiocrine and leukocyte-adhesion functions [3, 6, 61]. Moving towards mid-lobular zone 2) and peri-central regions (zone 3), interactions between perisinusoidal stroma and macrophages, including both resident kupffer and monocyte-derived macrophages, emerged as an increasingly prominent signal (**Fig. S3b,e**). Previous literature [62–64] suggests that hepatic stellate cells (HSCs), a key perisinusoidal stroma population, can be activated by Kupffer cells and monocyte-derived macrophages to produce extracellular matrix and pro-fibrotic factors, which further activates HSCs, shaping scar formation and fibrosis progression in chronic liver diseases. Overall, this spatial modulation of interaction partners along the PV–CV gradient is coherently recapitulated by LYNX, reflecting the layered hepatic microenvironmental organization linked to distinct multicellular communication states across the liver lobule.

### LYNX delineates co-existing DCIS and invasive tumor microenvironments in breast cancer

We next asked whether LYNX could uncover spatially organized cellular ecosystems in a more complex tissue setting with multiple coexisting malignant states. We applied LYNX to a primary breast cancer section containing both ductal carcinoma in situ (DCIS) and invasive tumor regions, two histopathologic states associated with distinct stromal, immune, and signaling microenvironments. As increasing evidence points to the local microenvironment as a key determinant of tumor progression, resolving the spatial organization of these coexisting ecosystems can reveal how different niches associated with distinct tumor states are simultaneously organized within the same tissue landscape [1, 10, 65]. This is particularly relevant in breast cancer, where DCIS, a pre-invasive lesion confined within the ductal epithelium, and invasive breast carcinoma represent biologically distinct disease stages, and multi-patient spatial studies have documented divergent stromal and immune microenvironments associated with each [66, 67]. We analyzed a tissue patch profiled with Xenium spatial transcriptomics containing 16,005 cells and 313-gene panel [19]. The paired histological images were used as the auxiliary modality informative of tissue architecture, and Xenium spatial transcriptomics as the primary modality. This region is particularly complex as it contains spatial transitions between immune-stromal periphery, two discrete DCIS cores, and surrounding invasive tumor regions within a single field of view (FOV) (**Fig. 3a**).

**Figure 3:**
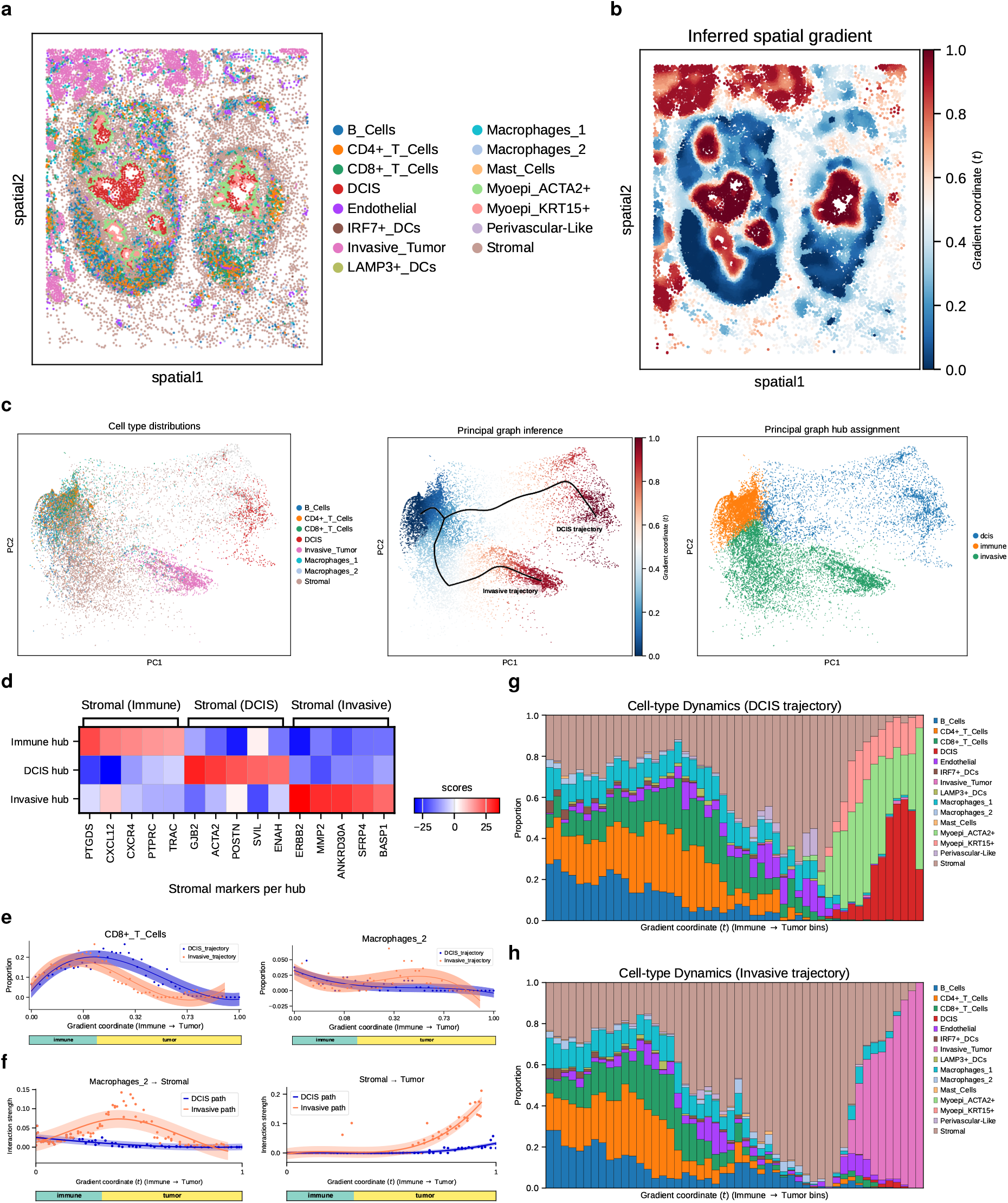
LYNX uncovers branching trajectories with distinct cellular dynamics across tumor progression states in human breast cancer. **a**, Spatial arrangement of cell states in the breast cancer replicate section containing both DCIS and invasive tumor cores [19]; DCIS: ductal carcinoma in situ. **b**, Inferred gradient coordinate (*t*) projected onto the spatial domain. **c**, From left to right: spatial distributions of immune, stromal and tumor states (**left**); Principal graph inference reveals a branching topology from the LYNX embedding, representing distinct trajectory paths from the shared immune-enriched “root” hub to different tumor branches (**middle**); discrete hub assignment with each cell mapped to its closest principal tree projection (**right, Methods**). **d**, Stromal markers differentially enriched with root, DCIS, and invasive hubs, identified based on hub assignments (**Methods**). **e-f**, Representative cell-type dynamics **(e)** and graded shifts of cell-cell interaction strengths **(f)** along each trajectory sorted by the inferred gradient coordinates; dots: observation; solid line: spline regression means; shaded areas: ± 1 s.d. **g-h**, Summary of cell-type dynamics along the DCIS trajectory (**g**) and the invasive trajectory (**h**). The inferred gradient coordinate is smoothed into 50 equal-width bins for dynamic computations (**Methods**).

Previous analysis of this tissue patch suggested a linear spatial gradient from the inner to outer regions, placing DCIS and invasive tumor at opposing ends of a single trajectory, with immune and stromal cells forming an intermediate continuum [50]. In contrast, LYNX revealed a branched trajectory topology from the multi-modal representation with a more flexible principal tree inference (see **Methods**) while faithfully reconstructing the primary transcriptomic profiles (Pearson’s *r* = 0.763; **Fig. S4a**). The inferred gradient maps coherently onto tissue space, with the immune-rich peripheral regions at the shared origin (*t* = 0), and the spatial coordinates extending outward along two branches that terminate in the DCIS and invasive tumor cores, respectively (**Fig. 3b,c, Fig. S4b**). This branching topology is corroborated at the phenotype level by continuous proportion dynamics along both branches. The shared origin is predominantly occupied by B cells, CD4^+^ and CD8^+^ T cells (**Fig. 3c**, left). Along both branches, immune abundance decreases and stromal populations become more prominent; the DCIS-associated branch terminates near myoepithelial and DCIS tumor-enriched regions, whereas the invasive-associated branch terminates near invasive tumor-enriched regions (**Fig. 3g,h**). By contrast, the previously reported linear spatial trajectory (invasive - ‘stromal - ‘immune - DCIS) yields fluctuating proportion dynamics, with stromal cells peaking in both the stromal and DCIS domain periphery [50].

To further examine our polarized spatial gradient pattern at the molecular resolution, we further assigned each cell to its nearest principal tree node in the embedding space, partitioning the continuous gradient into three contextual hubs, namely “immune”, “DCIS”, and “invasive” (**Fig. 3c**, right), and identified compartment-specific signatures along each hub for the underclassified stromal cells. Particularly, each of the three stromal contextual states carries a distinct transcriptional program (**Fig. 3d, Fig. S4c**) with a distinct layer separation around each hub (**Fig. S4d-e**). DCIS-adjacent stromal cells are enriched for *GJB2, POSTN*, and *ACTA2*, consistent with periductal desmoplastic activation; remodeling of the periductal extracellular matrix by stromal cells has been linked to DCIS encapsulation and progression risk [68, 69]. Invasive-adjacent stromal cells are instead marked by *ERBB2, MMP2* and *SFRP4*, consistent with a pro-invasive matrix-remodeling stromal state and spatial proximity to *HER2*-positive tumor compartments [19, 65]. Immune-adjacent stromal cells express immune-interface markers *CXCL12, CXCR4*, and *PTPRC*, suggesting chemokine-mediated immune retention at the tumor periphery. These trajectory-resolved states are further supported by spatially enriched cancer-associated fibroblasts (CAF) and Epithelial-mesenchymal transition (EMT) signatures along the invasive branch (**Fig. S4f-g**). Such enrichment of CAF programs in the invasive compartment is consistent with multi-patient spatial profiling linking cancer-associated fibroblast accumulation and collagen remodeling to invasive, but not in-situ, breast tumor stroma [66].

Across the inferred spatial gradient, CD8^+^ T cells and M1-like macrophages are more aggregated in the DCIS-proximal niche (*p* = 5.19 × 10^−3^, *p* = 2.59 × 10^−2^ respectively, 1-sided paired *t*-test) while M2-like Macrophages are enriched along the invasive path instead (*p* = 3.56 × 10^−3^); (**Fig. 3e**, see **Methods**), consistent with previous evidence of differential immune tumor-associated macrophage infiltration patterns between in-situ and invasive breast cancer stages [66, 69, 70], supporting the hypothesis of a more M2-associated immunosuppressive microenvironment in invasive tumor [65].

Beyond phenotype dynamics along the branching immune-stromal-tumor trajectories, LYNX also recovers key insights from cell-cell interactions. At the global level, interactions between lymphocytes to macrophages and endothelial cells shows strong signal (**Fig. S5a,b**). Moving locally towards DCIS and invasive tumor cores, we identify more prominent crosstalks between M2-like macrophages, stromal cells and tumor cells in the invasive hub compared to the DCIS hub (M2-like macrophage → stromal: *p* = 2.87 × 10^−6^; stromal → tumor: *p* = 7.80 × 10^−5^; M2-like macrophage → tumor: *p* = 3.31 × 10^−3^, 1-sided paired *t*-test) (**Fig. 3f, Fig. S5c**). These findings suggest that the spatially distinct organization of the invasive hub is reflected in coordinated remodeling of both stromal and immune compartments, including preferential enrichment of M2-like macrophage states and enhanced tumor-macrophage communication. This resonates with observations from Janesick et al. [19], where DCIS encapsulated by myoepithelial cell layers physically restricts tumor-associated macrophage infiltration (**Fig. 3a**). An independent study with multi-patient spatial proteomics profiling further shows that stromal cells are predominantly co-localized with tumor cells in invasive regions, consistent with a spatially compartmentalized tumor microenvironment across DCIS and invasive carcinoma [66].

To further study spatial communication flows, we applied COMMOT [33] to canonical, highly expressed receptor and ligand genes in the Xenium panel (**Fig. S6a,b**). CXCL12, enriched in stromal cells and macrophages, directs spatial flows toward peripheral CD4^+^ T cells (**Fig. S6a**), with polarity pointing outward from the stromal zone toward the immune periphery, consistent with a chemokine-retention role at the tumor–immune interface rather than passive spatial segregation (**Fig. S6b**). By contrast, IGF1 which is enriched in stromal and M2 macrophage regions showed spatial coupling with ERBB2-expressing tumor compartments (where ERBB2 encodes HER2). IGF1-ERBB2 flows converge inward toward both DCIS and invasive cores simultaneously, producing a sink-like pattern at each branch end (**Fig. S6b**), corroborating the spatially polarized stromal-to-tumor signal organization along the inferred gradient. Hub-level quantification further shows that IGF1-ERBB2 spatial coupling is selectively concentrated in the invasive niche relative to DCIS, consistent with the stronger macrophage-invasive tumor crosstalk identified above.

Collectively, the branching spatial gradient, hub-resolved stromal subtype partitioning, and cell-cell communication results together demonstrate LYNX’s capacity to dissect co-existing, spatially distinct tumor microenvironments within a heterogeneous tissue field, without presupposing a temporal ordering between disease states.

### LYNX recapitulates the cortico-medullary axis from spot-level mouse thymus data

To further demonstrate LYNX’s generalizability across tissue types and measurement platforms, we applied it to a paired spatial Stereo-CITE-seq dataset from mouse thymus [71], combining Stereo-seq transcriptomics (sequencing-based, transcriptome-wide) as the primary modality with CITE-seq proteomics as the auxiliary. Unlike Xenium, probe-based Stereo-seq offers full transcriptome coverage at the cost of lower spatial resolution at spot-level, while CITE-seq protein profiles are susceptible to signal degradation with limited 51-plex panels in this dataset.

The thymus organizes T cell maturation along a well-defined cortico-medullary axis (CMA): cortical thymic epithelial cells (cTECs) mediate positive selection in the outer cortex while medullary TECs (mTECs) enforce central tolerance in the medullary core, guiding thymocyte precursors from cortex to medulla during development [26, 71]. We adopted a recent semi-automated ground-truth CMA coordinate and zone annotations, including Capsule, Cortex, Cortico-Medullary Junction (CMJ) and Medulla, from [26] as the evaluation reference (**Fig. 4a**). This spatial organization provides a natural system for studying how coordinated molecular programs emerge along a continuous developmental trajectory within tissue space.

**Figure 4:**
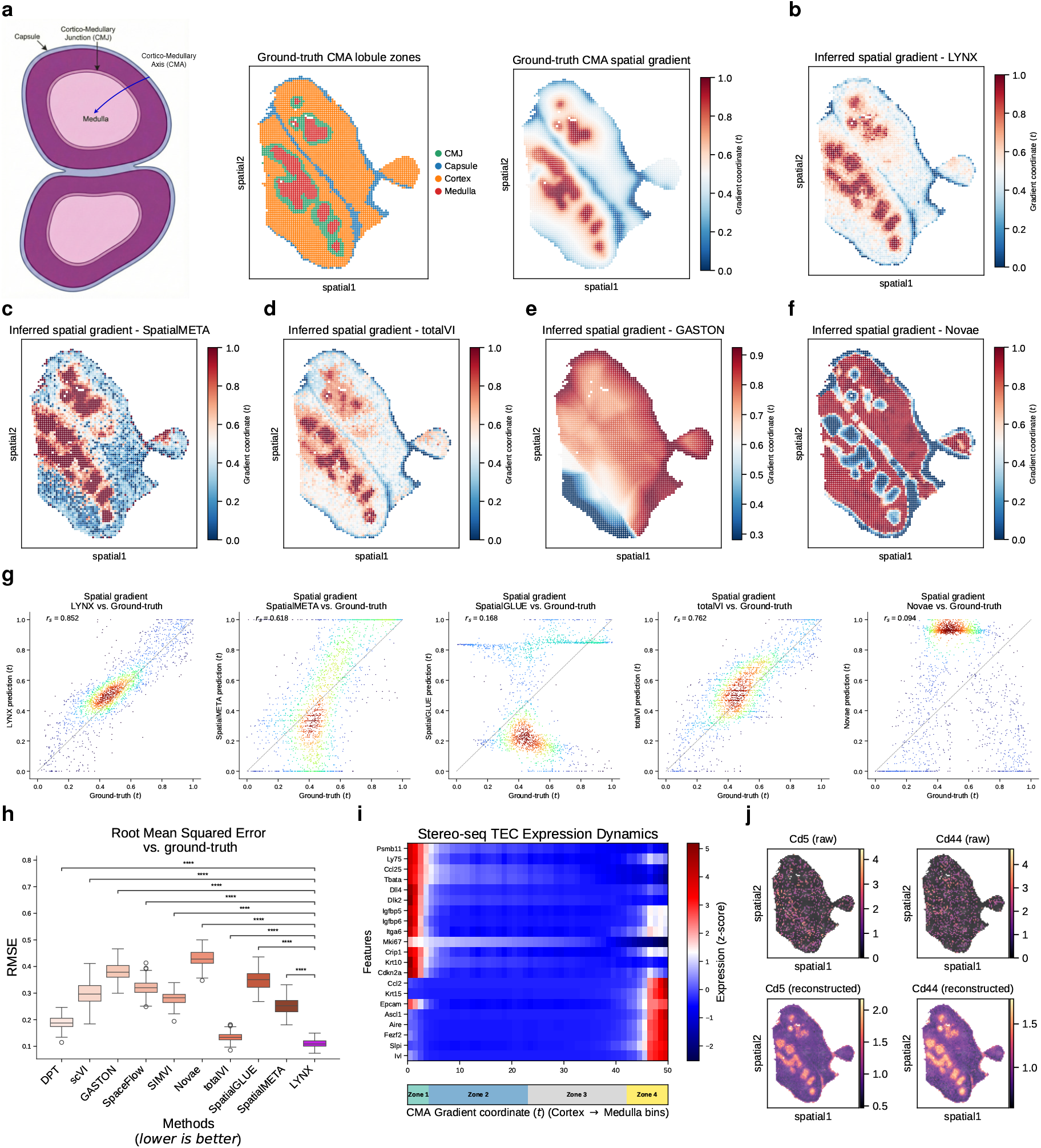
LYNX recapitulates the cortico-medullary axis (CMA) and T cell migration trajectories in the mouse thymus. **a**, Diagram of the thymus anatomical structure along the CMA (**left**) [26]; spatial visualizations of the annotated ground-truth thymus lobule layers (**middle**) and continuous CMA coordinate *t* (**right**). **b**–**f**, Spatial distributions of inferred gradients from LYNX (**b**) and other multi-omics integration and spatial gradient methods: spatialMETA (**c**), totalVI (**d**), GASTON (**e**) and Novae (**f**). **g-h**, Quantitative benchmarking of inferred spatial gradient coordinate from LYNX and competing methods against ground-truth CMA: Spearman Correlation Coefficient **(g)**; root-mean-squared error (RMSE) **(h)**; boxplots indicate the median (center lines), interquartile range (hinges) and 1.5× interquartile range (whiskers); statistical significance was computed with a one-sided t-test; \*\*\*\**p <*1e-4. **i**, Stereo-seq RNA expression dynamics of thymic epithelial cell (TEC) markers along LYNX gradient coordinate. **j**, Example comparison of marker gene expressions between raw (top) and LYNX denoised reconstruction (bottom). Log-normalize expression counts are reported for visualization.

LYNX accurately recovered the CMA spatial gradient, producing the closest agreement with independently annotated spatial coordinates among all benchmarked methods (significantly higher Spearman correlation and lower RMSE; **Fig. 4b,g,h**). Methods that performed competitively on the liver dataset, including Novae and spatialMETA, were substantially outperformed here (**Fig. 4c–f**), indicating that LYNX’s gains reflect cross-dataset generalizability rather than delicate pre-training and tissue-specific tuning. For discrete zone recovery, although the anatomically narrow CMJ posed a difficult ground-truth annotation, LYNX attained the best NMI and AMI scores, accurately delineating the structural differences between outer capsule, cortex and medulla (**Fig. S7a,b**). Gradient spatial smoothness, revealed by Moran’s I, was also among the highest except from GASTON, whose over-smoothed predictions inflate this metric at the cost of structural accuracy (**Fig. S7c**).

Having established accurate recovery of thymic spatial organization, we next examined the molecular programs associated with the inferred developmental axis. The inferred spatial axis revealed coordinated transcriptional programs associated with thymic maturation and epithelial specialization[26]: cortex-enriched cTEC markers (*Psmb11, Ly75* and *Ccl25*) concentrate in capsule and cortex region, while medullary mTEC markers (*Aire, Fezf2, Krt15, Epcam*, and *Ascl1*) rise progressively in zones 3–4 corresponding to CMJ and medulla (**Fig. 4i**).

In contrast to the relatively discrete epithelial organization, macrophage populations exhibited gradual spatial transitions across the axis, suggesting continuous remodeling of immune-supportive niches during thymocyte maturation(**Fig. S7e**): cortex-enriched markers *Cd163, Hpgd*, and *Cx3cr1* decline toward the medullary core, while *Timd4, Cd63*, and *Zmynd15* aggregate towards the end of the gradient, which align well with observations from Liao et al. [71]. CITE-seq protein dynamics provides further evidence of the spatial organization: *CD4* and *CD8a* protein levels decrease progressively from cortex to medulla (**Fig. S7f**), consistent with the transition from cortical double-positive (DP) thymocytes toward medullary single-positive populations [72]; CD169 peaks at the CMJ, indicative of its sinusoidal macrophage localization at the cortico-medullary boundary [71].

Finally, multimodal integration uncovered spatial programs that were difficult to resolve from either modality individually. Two established markers of mature medullary T cells, *Cd5* and *Cd44*, are nearly undetectable in both the raw CITE-seq protein and Stereo-seq RNA signals (**Fig. 4j**). Despite the weak signal in both modalities, LYNX recovered spatially coherent expression patterns that localized specifically to the medulla. Unlike conventional multimodal integration frameworks that jointly reconstruct all modalities, LYNX leverages the auxiliary modality to condition a shared latent representation while concentrating generative capacity on the primary modality. This design allows complementary spatial information to be transferred across modalities while mitigating propagation of modality-specific noise, thereby enhancing recovery of biologically interpretable spatial structure. Additional markers with strong region-specific aggregations are also recovered, such as *Mi67* (cortex) and *Serpinb2* (medulla) (**Fig. S7d**).

These results suggest that tissue architecture itself can serve as an informative constraint for recovering biologically relevant molecular programs from noisy multimodal measurements. This also demonstrates that LYNX’s hierarchical design enables meaningful cross-modal signal recovery even when the secondary modality is of markedly inferior quality, strengthening a practical solution for paired multi-omics datasets where modality quality is often unequal.

### LYNX enables 3D volumetric gradient inference from multimodal spatial atlases

Having shown that LYNX recovers continuous spatial gradients across 2D multimodal tissue sections in liver, breast cancer, and thymus, we next asked whether the same framework could extend beyond single-section analysis to reconstruct spatial organization across tissue depth. Returning to the liver portal-to-central vein (PV–CV) axis that anchored the first application, we asked whether the 2D liver gradient recovered from a single Xenium section reflects genuine three-dimensional tissue organization or is susceptible to section-level sampling bias, a concern well-documented in 3D spatial atlases of cancer where individual cross-sections have been shown to miss spatial state transitions and immune interaction patterns only apparent across tissue depth [73, 74]. Prior volumetric reconstructions of the hepatic lobule have been largely restricted to single CV–PV neighborhoods resolved by high-resolution intravital or confocal microscopy, capturing tissue volumes orders of magnitude smaller than a whole-slide field of view and relying predominantly on murine models [48, 75].

Most existing spatial gradient, spatial domain, and multimodal integration methods are formulated for single 2D sections; when applied to serial sections, they typically require independent per-section analysis, matched resolutions, or post hoc alignment of embeddings rather than joint volumetric multimodal inference. We therefore extended LYNX to a volumetric multi-modal setting using seven consecutive tissue sections from the same patient used in the 2D study, spanning *>*800 µm of tissue depth, with each block comprising a DESI^+^/Xenium/DESI^−^ sandwich triplet at 10 µm inter-section spacing (**Fig. 5a**).

**Figure 5:**
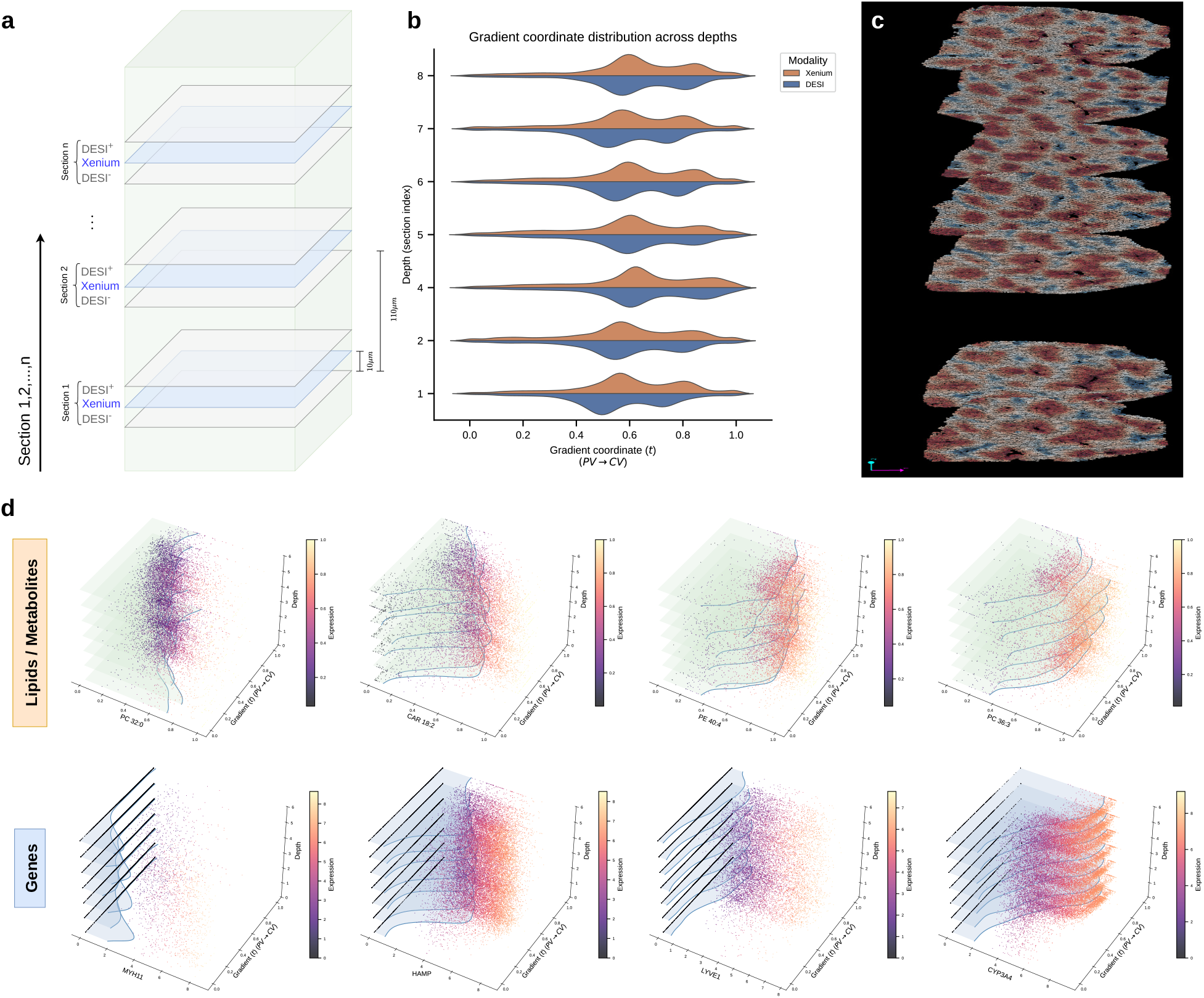
LYNX reconstructs the full 3D porto-central axis in liver. **a**, Diagram of tissue retrieval for 3D atlas staining; Adjacent section groups were collected at 110 µm intervals; within each group, the central slice was used for the Xenium spatial transcriptomics experiment, whereas lipids & metabolites with positive / negative ions were acquired on the two flanking slices 10 µm apart (DESI^+^/DESI^−^). **b**, Distributions of the inferred porto-central gradient coordinate across individual tissue section blocks. **c**, Side-view (*z*-axis ↑) of full 3D visualization of the portal-central spatial gradient inferred by LYNX. **d**, Visualizations of canonical markers with zonal distributions. **Top**: DESI markers; **Bottom**: Xenium markers. Dots represent DESI intensities (scaled to [0, 1]) and Xenium expressions (log-normalized) across all cells; Shaded areas under the curves represent fitted kernel-density estimates per section depth. Each marker’s expression is visualized across all seven sections (*x*-axis), with each cell colored by its inferred gradient coordinate *t* (*y*-axis) stacked by section depths (*z*-axis).

Integrating this z-stack required solving three hierarchical registration tasks: (i) iterative cross-section Xenium co-registration, where each section is aligned to a central fixed reference by affine transformation; (ii) intra-section DESI^+^/DESI^−^ channel alignment; and (iii) cross-modal Xenium-to-DESI alignment at each depth level, bridging the high spatial resolution gap between modalities. All three were handled through a unified anchor-based affine pipeline implemented in `napari-spatialdata` [76, 77], in which users iteratively annotate paired landmark points across modalities / sections via interactive visualization (see **Methods**). The registered seven-section stack was then passed to LYNX for multi-slice inference, with all Xenium–DESI section pairs trained jointly under a single shared model rather than analyzed independently. LYNX’s standard subgraph-partitioned training (see **Methods**) scaled naturally to this larger volumetric graph, after which LYNX inferred a common PV–CV gradient coordinate that could be projected across the 3D tissue landscape.

The 3D rendering of LYNX-inferred porto-central spatial gradient reveals that the characteristic lobular architecture, ranging from portal triad islands to pericentral territories, is structurally smooth and continuously reproduced across all seven sections without obvious depth-dependent distortions (**Fig. 5b,c**). To further quantify cross-depth gradient consistency, we examined gradient coordinate-resolved expressions stratified by section depth (**Fig. 5d**). Periportal marker *MYH11* and pericentral marker *CYP3A4* each display a flat, monotone expression distribution that overlaps across all seven z-levels without systematic drift, whereas *HAMP* and *LYVE1* present non-monotonic dynamics peaking at mid-zone regions. Similarly, our 3D DESI results show consistent portal-vein aggregation of *PC 32:0*, a saturated phosphatidylcholine with increased signals in human cirrhotic liver [58], whereas the unsaturated counterparts such as *PC 36:1* and *PC 36:3* are distributed towards central veins, consistent with our 2D results (**Fig. 2h, Fig. 5d**) and a previous MSI data from human liver [78].

Altogether, the concordant 2D and 3D results confirm that LYNX reconstructs a stable, cross-modal, coherent PV-CV gradient at both single-section and volumetric resolution, with transcriptomic and metabolomic zonation markers reproducing reliably across high depth of tissue blocks. In the future, fully characterizing hepatic zonation will ultimately require complementing these molecular readouts with biophysical measurements, such as bile flow direction and canalicular pressure, which currently remain accessible only through live intravital imaging followed by longitudinal computational fluid modeling [79].

## 3 Discussion

We introduced LYNX, a hierarchical generative framework for inferring continuous spatial gradients and gradient-resolved cell–cell interactions from paired multi-modal spatial data. Our work is motivated by a central property of tissue organization: molecular programs, cellular states, and intercellular interactions often vary continuously across space rather than being confined to discrete compartments. While existing spatial methods have advanced discrete domain identification [23–25], representation learning [28, 46, 52], and cell-cell communication analysis [33–35], these capabilities have largely developed independently, and the few methods that address spatial gradients do so within a single modality and without coupling to cellular interaction dynamics [50, 51]. LYNX brings them together in a unified interpretable framework that learns continuous spatial organization from heterogeneous multimodal measurements and relates this organization to both molecular dynamics and changes in cellular interaction networks.

A key design principle of LYNX is an asymmetric conditional generative process in which the coarser or lower-throughput auxiliary modality shapes a flexible niche-level latent prior without being explicitly reconstructed, concentrating generative capacity on the higher-fidelity primary readout. This design addresses two pervasive challenges in multimodal spatial profiling: substantial disparities in spatial resolution and throughput across assays, and heterogeneous modality-specific noise characteristics. By conditioning latent structure on auxiliary spatial information while reconstructing only the primary modality, LYNX enables integration of modalities with non-overlapping feature spaces and markedly different resolutions without requiring feature matching or resolution harmonization [27–29]. We further justified LYNX’s multimodal design via rigorous ablation studies, demonstrating remarkable performance gains from hierarchical auxiliary conditioning, correct multi-modal alignment and integration on the liver and thymus applications (**Fig. S8**, see **Methods**). Benchmarked across transcriptomics–metabolomics, transcriptomics–proteomics, and transcriptomics–histology pairings against a diverse panel of spatial domain, multi-modal integration, and gradient inference baselines (many of which are themselves graph neural network-based [27, 52, 53]), LYNX consistently achieved superior gradient recovery, spatial clustering accuracy, and runtime efficiency.

The biological applications demonstrate that continuous gradient modeling uncovers tissue organization that discrete domain approaches would obscure. In human liver, the reconstructed PV-CV axis confirmed established metabolic zonation patterns [2, 8] and revealed zone-dependent remodeling of intercellular signaling, including LSEC-dominated communication in periportal zones and an emerging perisinusoidal stroma–hepatocyte interaction axis toward the central vein, alongside pericentral *LGR5* expression consistent with active Wnt/*β*-catenin amplification [8, 59]. In human breast cancer, LYNX resolved spatially branching trajectories across DCIS and invasive compartments, partitioning the stromal population into three contextually distinct states and uncovering differential macrophage–tumor interaction strengths that track with invasion status. In mouse thymus, LYNX faithfully recovered the cortico-medullary axis alongside the expected spatial segregation of TEC subtypes and macrophage niches [26, 71, 72], while additionally demonstrating that a severely degraded CITE-seq auxiliary modality could be rehabilitated through the hierarchical conditioning design. Extending to a seven-section, *>*800 µm volumetric stack from the same liver specimen confirmed that these 2D gradient inferences reflect consistent three-dimensional zonation biology rather than section-level sampling artifacts.

Beyond the individual biological applications, our results highlight a broader conceptual perspective for spatial omics analysis. LYNX models tissues as continuous spatial landscapes over which molecular programs, cellular phenotypes, and interaction networks evolve. This representation enables comparisons across modalities, spatial scales, and biological systems within a shared latent framework. Importantly, the same formulation naturally extends from 2D to 3D tissue multimodal datasets, providing a path toward integrated molecular atlases that preserve both spatial continuity and multimodal context.

Several limitations suggest directions for future work. First, cell–cell interaction inference is based on latent interaction weights rather than explicit ligand–receptor modeling, limiting mechanistic interpretation. Although *LGR5* emerged as a pericentral Wnt marker, its primary ligand *RSPO3* was absent from the Xenium panel, and integration with more comprehensive ligand–receptor databases and gene panels will strengthen pathway-level interpretation [33, 80, 81]. Second, LYNX reconstructs molecular organization but not biophysical tissue properties such as bile flow or canalicular architecture, which require complementary imaging and physical modeling [79]. Third, although the volumetric application demonstrates the feasibility of joint 3D multimodal inference, denser z-axis sampling and larger tissue volumes will be needed for complete organ-scale reconstruction.

Looking forward, the modality-agnostic architecture of LYNX naturally extends to emerging paired spatial profiling technologies and larger volumetric atlases. As multimodal spatial measurements continue to diversify, LYNX provides a unified framework for studying how molecular programs and cellular interactions evolve across continuous tissue landscapes.

**Figure S1:**
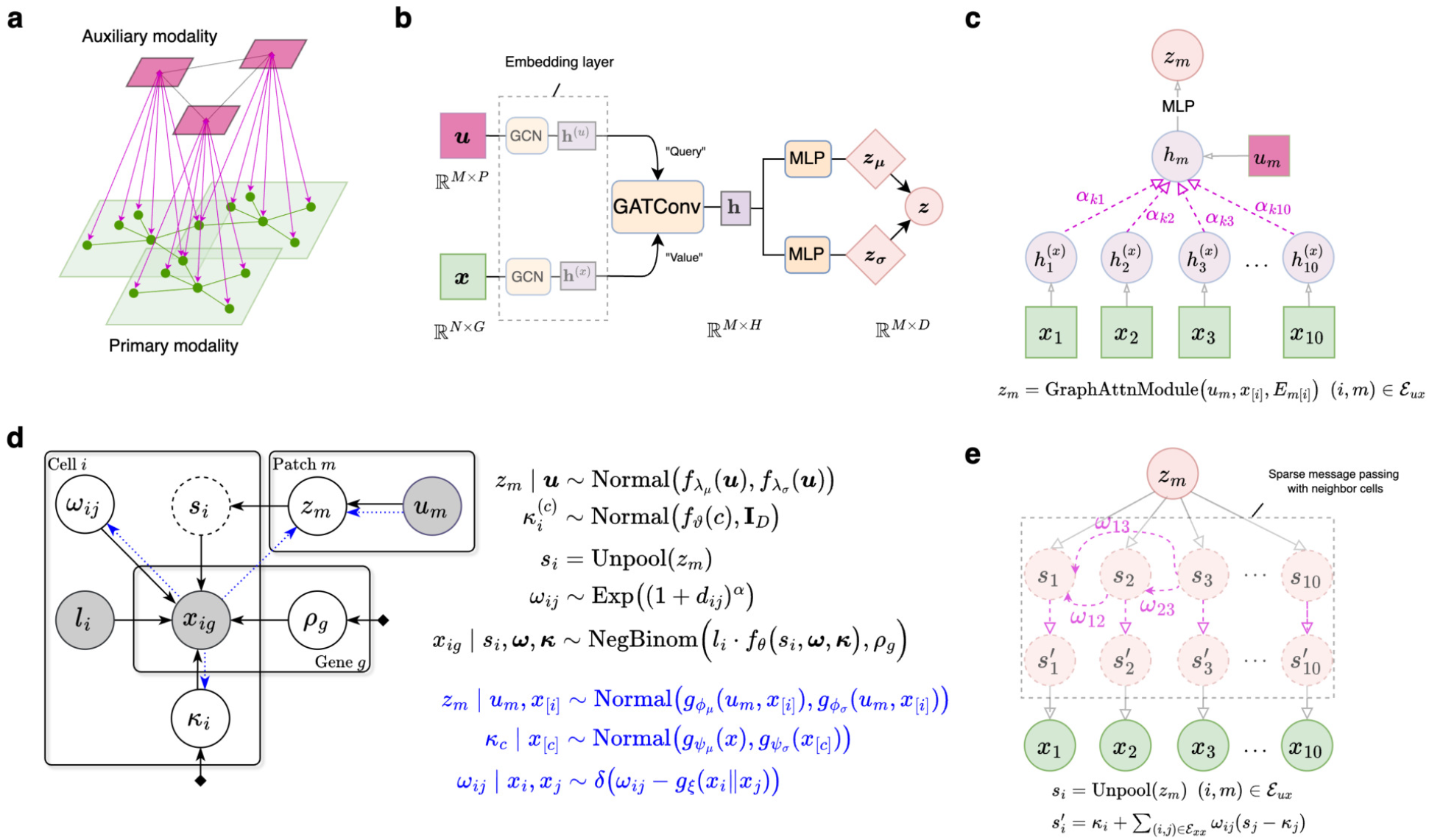
LYNX model and architecture specifications. **a**, Schematic diagram of multi-modal spatial heterogeneous graph. *Nodes*: primary modality samples (e.g. cells), auxiliary modality samples (e.g. spots, image pixels / patches); *Edges*: auxiliary-auxiliary, primary-primary, auxiliary-primary. **b-c**, LYNX encoder architecture diagrams. Full variational inference from paired spatial observations to the joint niche embedding (**b**); Multi-modal integration with graph attention module highlighted through an example subgraph: primary nodes (*x*_1_, …, *x*_10_) are spatially aligned to auxiliary node *u*_*m*_. **d**, Full graphical plate model of the LYNX generative path (decoder) and variational inference path (encoder; shown in blue). **e**. LYNX decoder with the niche-cell unpooling illustrated through an example subgraph.

**Figure S2:**
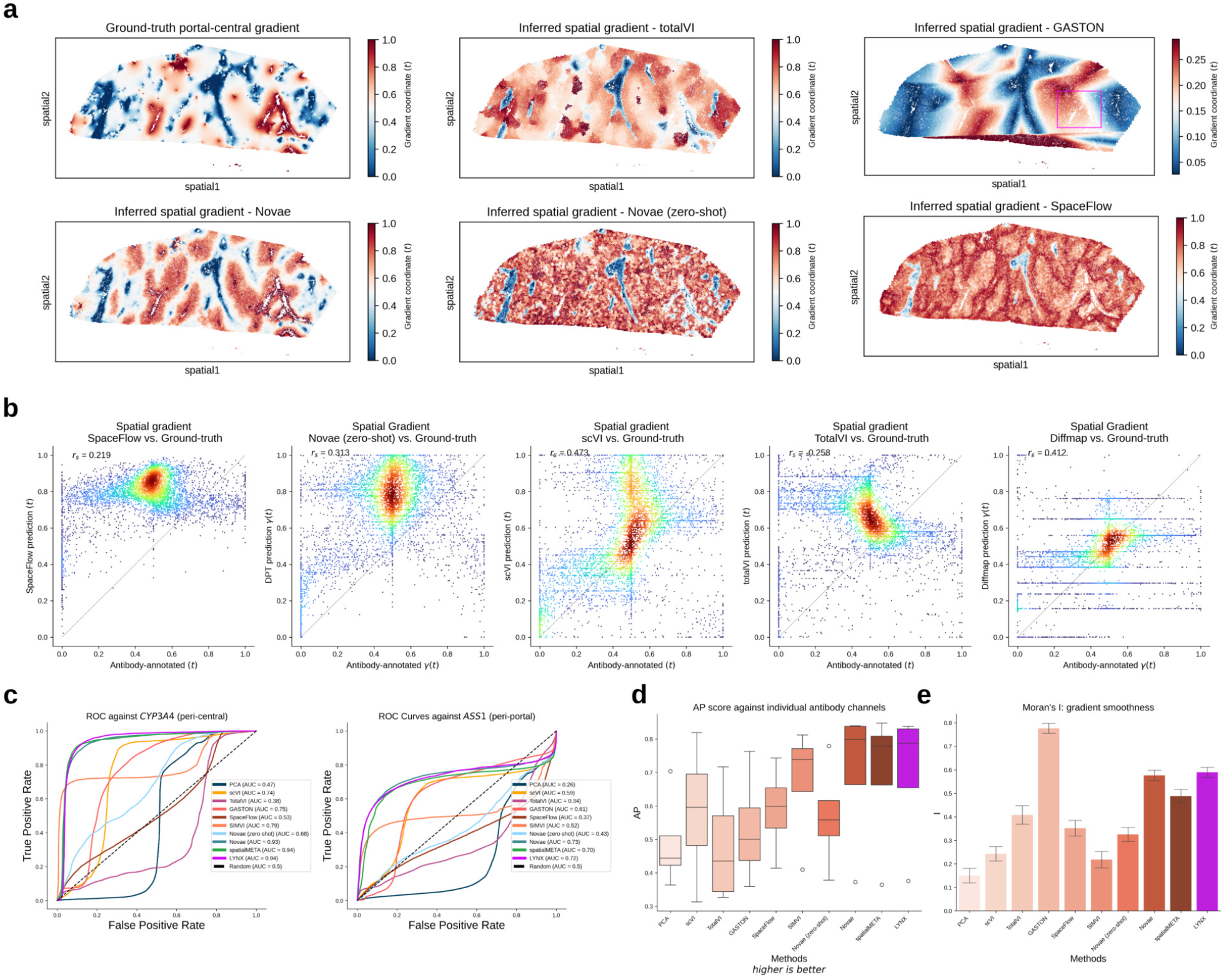
Benchmark summary on paired Xenium-DESI human liver sample. **a**, Spatial gradients of the porto-central axis from antibody generated ground-truth proxy (**Methods**) and other inference methods. **b**, Spearman correlation between the inferred methods and the ground-truth gradient. **c**, Receiver Operating Characteristic (ROC) curve between LYNX and other methods against held-out antibody staining channels. **Left**: pericentral (PC) marker, **Right**: periportal (PP) marker. **d**, Average Precision (AP) score across benchmarked methods against individual antibody staining channels. **e**, Spatial smoothness across all the inferred gradients. In panels **d,e**, boxplots indicate the median (center lines), interquartile range (hinges) and 1.5× interquartile range (whiskers).

**Figure S3:**
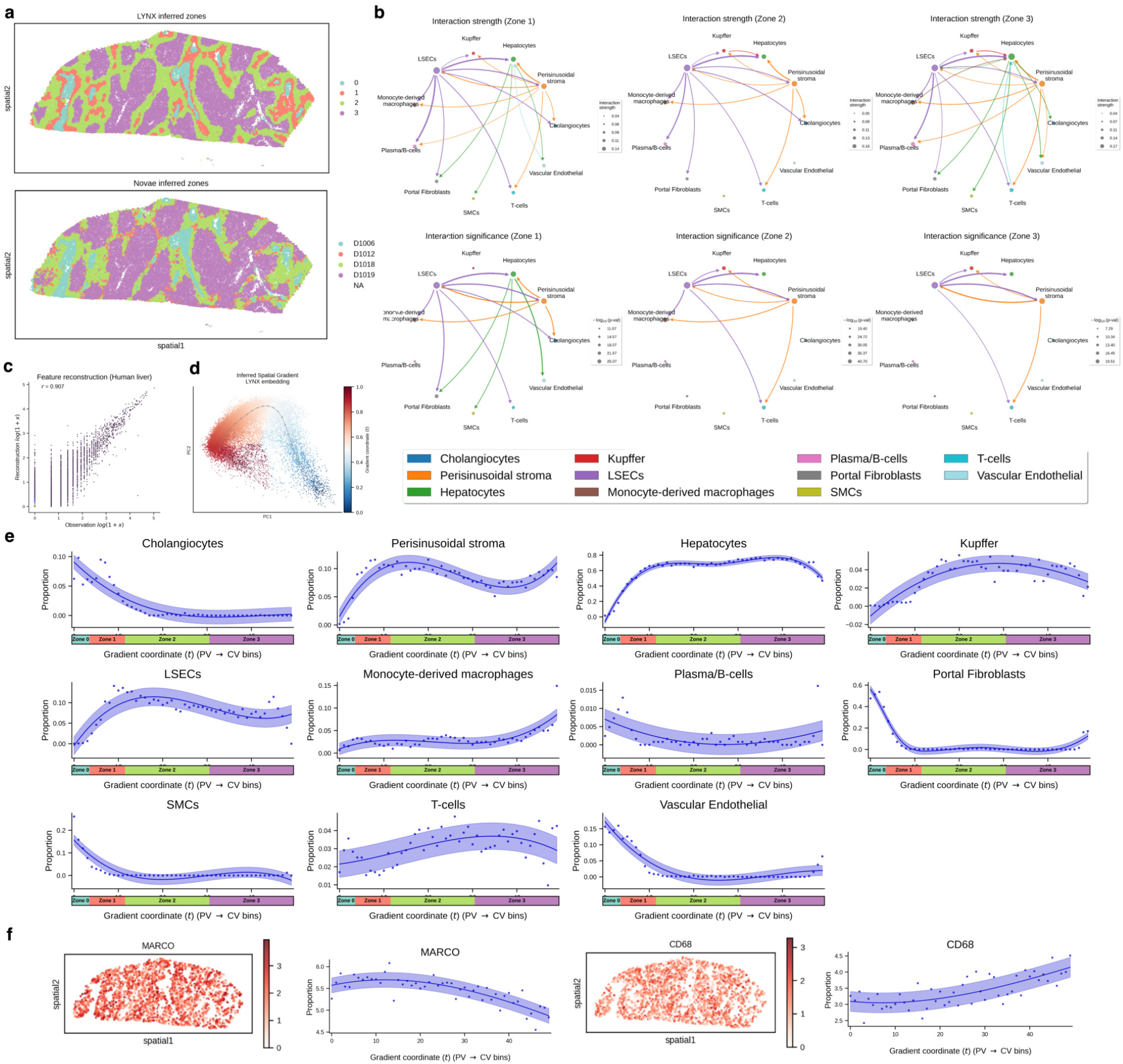
Downstream analysis on paired Xenium-DESI human liver sample. **a**, Spatial zonation comparison along the predicted porto-central axis. **Top**: LYNX predictions, with contiguous boundaries from the portal triad-proximal zone (zone 0) to metabolic zones (zones 1–3) towards central-vein. **Bottom**: Novae zone assignments (fine-tuned) on the same tissue section, exhibiting spatially scattered and fragmented boundaries in metabolic zones 2 and 3. **b**, Inferred cell-cell interaction strength (top) and significance (bottom) across the three canonical metabolic zones (zones 1–3; periportal, mid-lobular, and pericentral). Interaction strength summarizes the raw LYNX predictions, whereas significance reports the − log_10_(p-val) tested against the cell-type abundance baselines via paired one-sided t-test (**Methods**). **c**, LYNX feature expression reconstruction of the primary modality (Xenium). **d**, Principal graph inference on top of the corresponding latent embedding visualized with PCA. **e**, Individual phenotype dynamics along the LYNX inferred porto-central axis. **f**. Expression distributions of representative macrophage markers (MARCO & CD68) along the spatial domain and LYNX inferred porto-central axis. **e-f**: dots: observation; solid line: spline regression means; shaded areas: ± 1 s.d.

**Figure S4:**
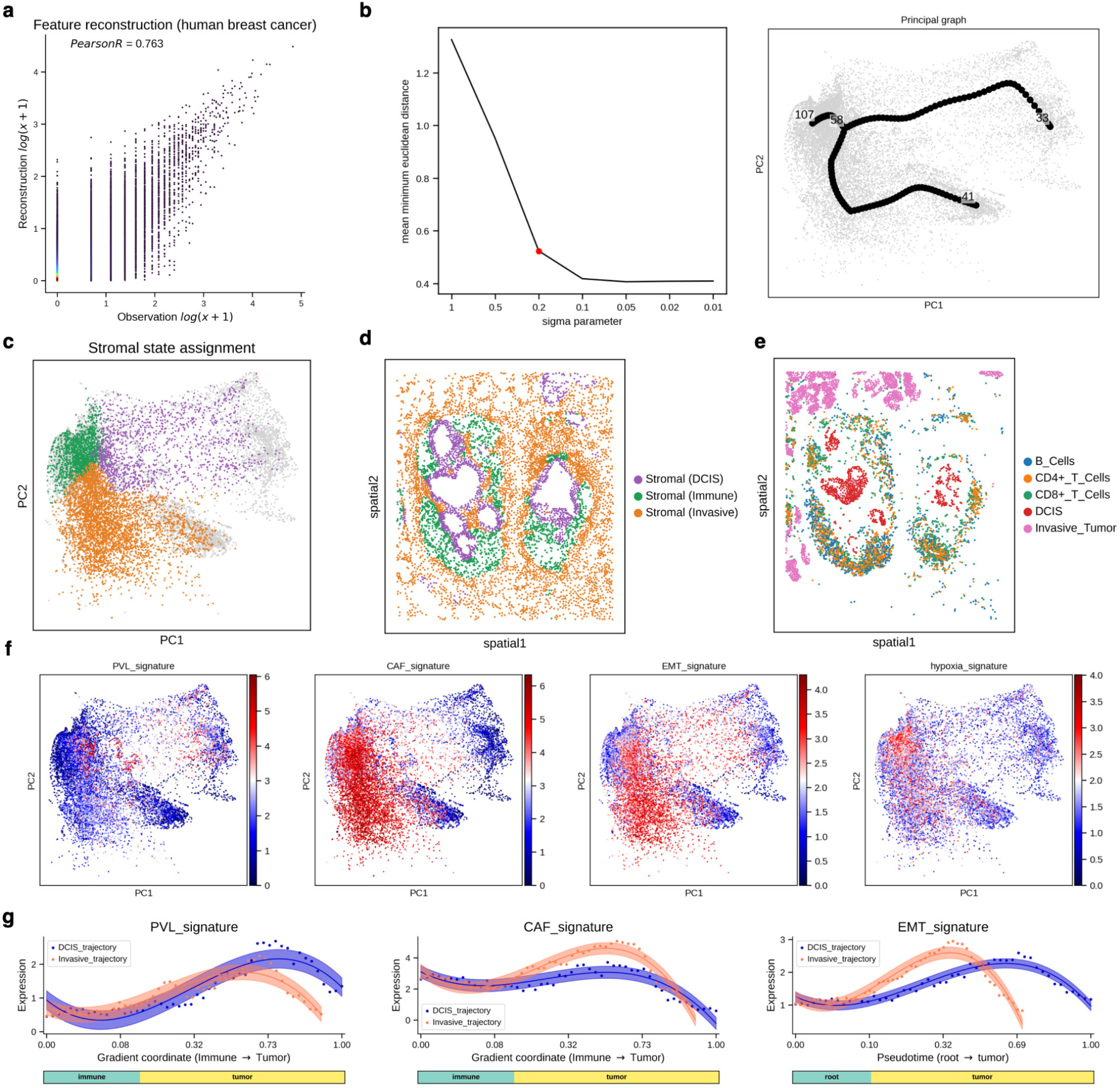
Downstream analysis on paired Xenium-histology human breast cancer sample. **a**, LYNX feature expression reconstruction of the primary modality (Xenium). **b**, Principal tree inference on top of the corresponding latent embedding. **Left**: hyperparameter search for the elastic principal tree; **Right**: PCA visualization. **c**, PCA visualization of the latent embedding colored with stromal state assignment. All the stromal cells are assigned to the closest hub (Immune, DCIS or Invasive) based on their embedding distance along the principal tree. **d-e**, Spatial distributions of stromal hub assignments **(d)**, immune and tumor states **(e). f**, PCA visualization of the latent embedding colored with Pathway signature expressions. **(g)**, Dynamics of pathway expressions along each trajectory sorted by the inferred gradient coordinates

**Figure S5:**
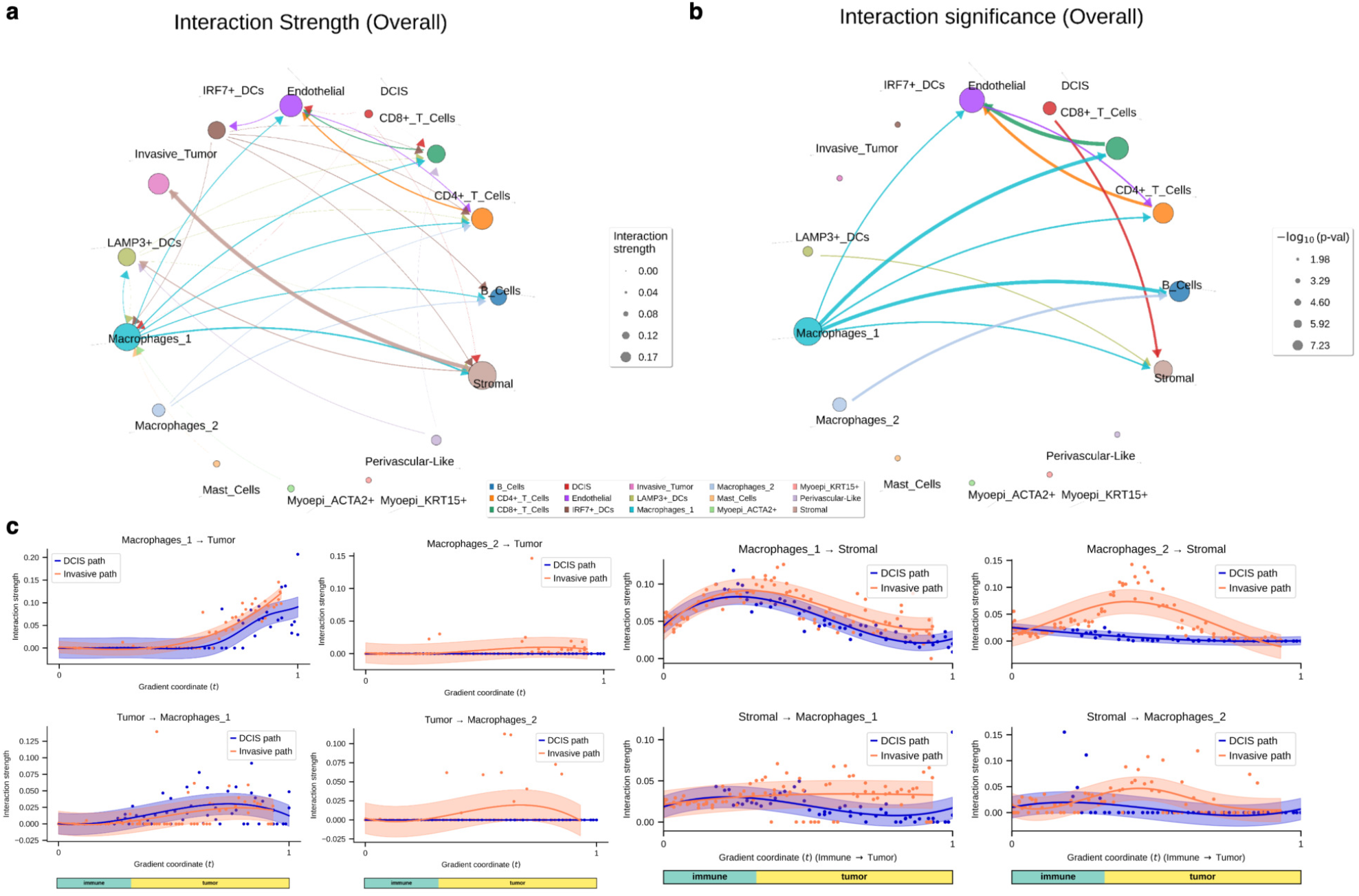
Cell-cell interaction analysis on paired Xenium-histology human breast cancer sample. **a-b**, Overall summary of cell-cell interaction strength (**a**) and significance over co-localization baselines (**b**) (**Methods**). **c**, Localized macrophage-tumor interaction strengths between macrophages, stromal and tumor cells along the inferred spatial gradient coordinate. dots: observation; solid line: spline regression means; shaded areas: ± 1 s.d..

**Figure S6:**
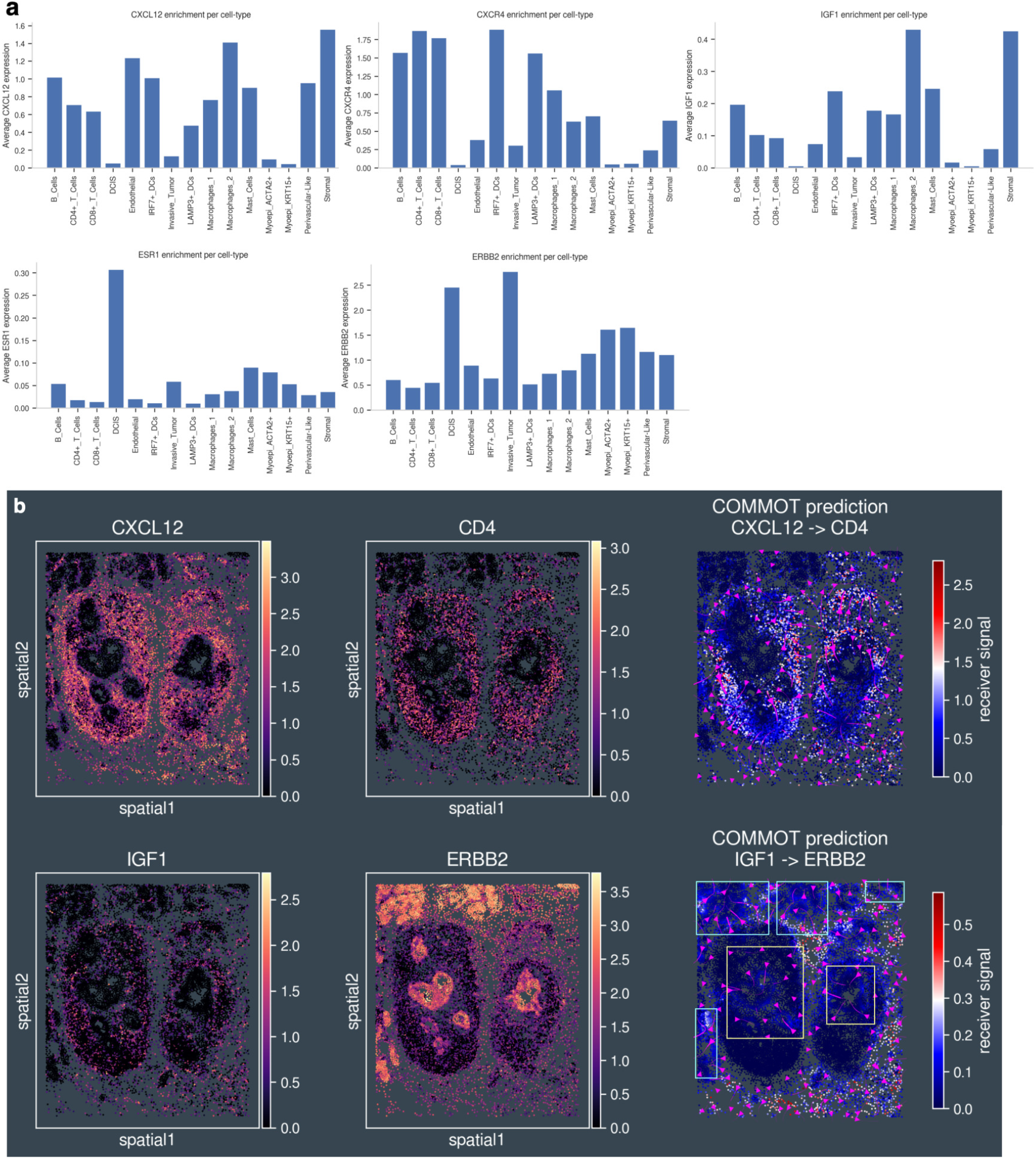
Validation of LYNX results with representative ligand-receptor pairs on human breast cancer. **a**, Representative ligand and receptors enriched in stromal, macrophages (*IGF1, CXCL12*), lymphocyte (*CXCR4*) and tumor states (*ERBB2*). **b**, Spatial distributions of representative ligand-receptor pairs and their signaling strengths from COMMOT [33]. **Top**: stromal/macrophage → T-cells, **Middle** and **Bottom**: stromal/macrophage → tumor; yellow bounding boxes: DCIS compartment; cyan bounding boxes: invasive compartment.

**Figure S7:**
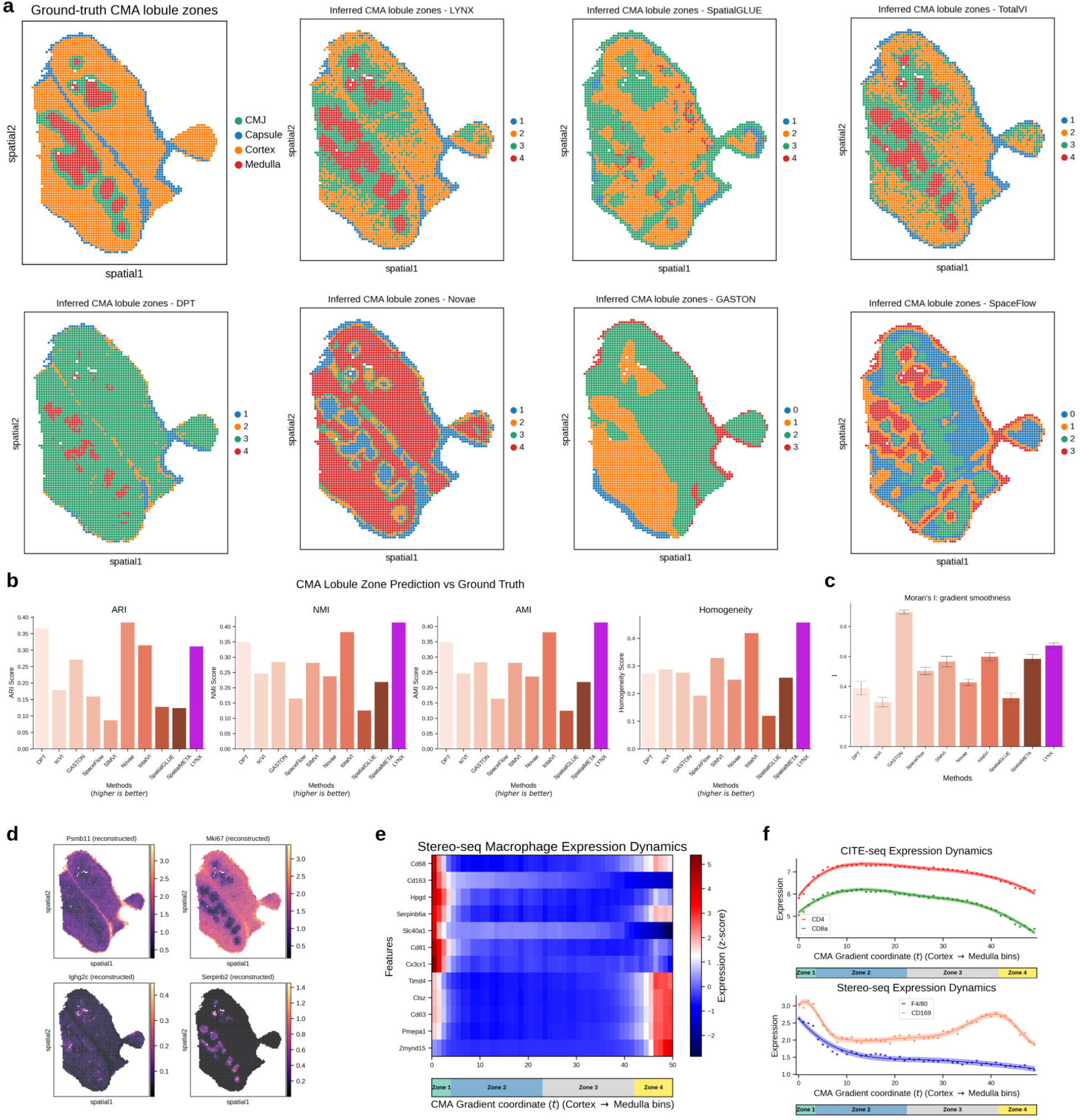
Benchmark and downstream analysis on Stereo-CITE-seq mouse thymus data. **a**, Spatial visualization of the annotated ground-truth CMA lobule zones and predictions from LYNX and competing methods. **b**, Clustering accuracy between prediction methods against the ground-truth. Left-to-right: Adjusted Rand Index (ARI), Normalized Mutual Information (NMI), Adjusted Mutual Information (AMI). **c**, Spatial smoothness across all the inferred gradients. **d**, Comparison of the raw RNA expressions (top) and LYNX reconstructions (bottom). **e**, Macrophage RNA-seq expressions sorted along the LYNX inferred CMA axis. **f**, Example T-cell (top) and macrophage (bottom) marker expressions along the LYNX inferred CMA axis. dots: observation; solid line: spline regression means; shaded areas: ± 1 s.d.

**Figure S8:**
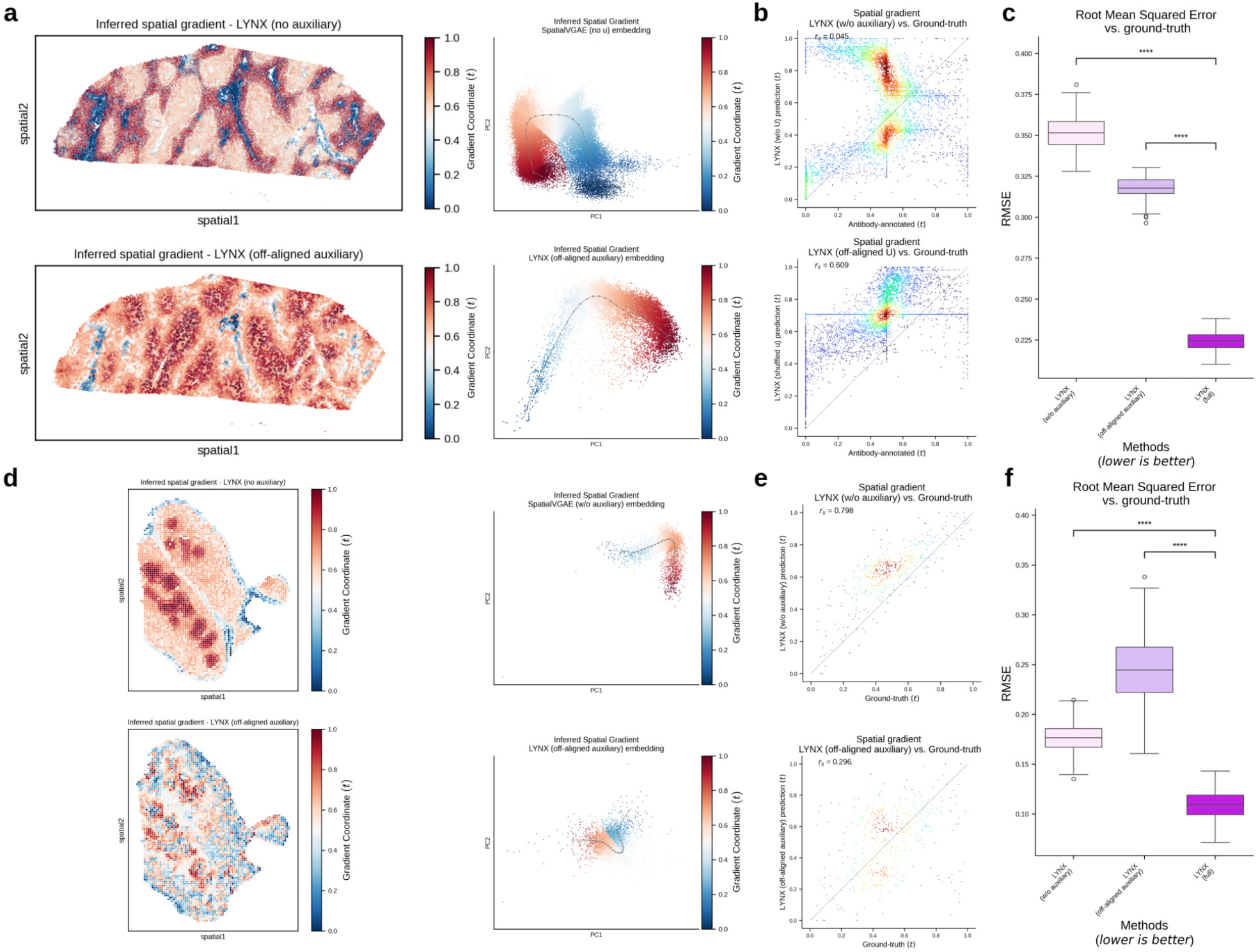
Ablation studies on the LYNX model design. **a-b**, Ablation study of reduced LYNX model on the human liver dataset. From left to right: visualizations of the predicted gradient coordinate on spatial latent domains (**a**) and scatterplot against the ground-truth gradient (**b**). **Top**: LYNX without the auxiliary modality. **Bottom**: LYNX with spatially off-aligned auxiliary-primary. **c**, RMSE comparisons of full vs. reduced LYNX architecture. **d-e**, Ablation study of reduced LYNX model on the mouse thymus dataset. From left to right: visualizations of the predicted gradient coordinate on spatial latent domains (**d**) and scatterplot against the ground-truth gradient (**e**). **Top**: LYNX without the auxiliary modality. **Bottom**: LYNX with the auxiliary modality. **f**, RMSE comparisons of full vs. reduced LYNX architecture In panels **c, f**. One-sided independent t-test were used for significance tests, **** *p <*1e-4; boxplots indicate the median (center lines), interquartile range (hinges) and 1.5× interquartile range (whiskers).

## Methods

### 1 The LYNX model

#### Overview

LYNX is a deep generative model that learns spatial gradients and cellular interaction dynamics through a joint latent manifold from paired spatial-omics data. Inspired by previous multi-modal integration methods, the core LYNX architecture extends upon a conditional VAE with a graph-based encoder and a linear decoder [82, 83]. LYNX takes a primary modality and a secondary, “auxiliary” modality as input, and reconstructs the primary observations conditioning on the auxiliary as output (**Fig. 1**). The paired inputs are allowed to have varying resolutions and throughput as long as they’re acquired from adjacent tissue slices.

#### Notations

Let 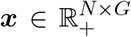 be the expression matrix of the primary modality and 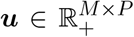 for the auxiliary modalities, where *N* and *M* could represent the number of cells and patches respectively. By convention, we use “high-resolution” for primary observation’s sample size and “low-resolution” for auxiliary interchangeably. To link spatial neighbors across datasets, a heterogeneous graph *G* = (*V, E*) is constructed with both *within*-modal and *cross*-modal edges (see **Methods: Multi-modal graph construction**). Let *z* ∈ ℝ^*M*×*D*^ and *s* ∈ ℝ^*N*×*D*^ represent latent niche representations with respect to the auxiliary and primary modality resolutions. We denote *γ* : [0, 1] → ℝ^*D*^ as the spatial trajectory mapping a normalized gradient coordinate *t* ∈ [0, 1] to LYNX latent embeddings. The following table summarizes the full notations:

| Notation | Definition |
| --- | --- |
| $\mathbf{x} \in \mathbb{R}_+^{N \times G}$ | primary modality expressions |
| $\mathbf{u} \in \mathbb{R}^{M \times P}$ | auxiliary modality expressions |
| $\mathbf{z} \in \mathbb{R}^{M \times D}$ | niche embedding (low-resolution latent) |
| $\mathbf{s} \in \mathbb{R}^{N \times D}$ | cell embedding (high-resolution latent) |
| $c_i \in \{1, \dots, C\}$ | discrete cluster or cell-type label for cell $i$ |
| $\boldsymbol{\kappa}_i \in \mathbb{R}^D$ | cell-level baseline offset with cluster-conditioned prior |
| $l_i \in \mathbb{N}_+$ | library size for cell $i \in \{1, \dots, N\}$ |
| $\omega_{ij} \in \mathbb{R}_+$ | cell-cell interaction strength from source $j$ to target $i$ |
| $\rho_g \in \mathbb{R}_+$ | inverse dispersion for gene $g \in \{1, \dots, G\}$ |
| $\gamma : [0, 1] \rightarrow \mathbb{R}^D$ | spatial trajectory |
| $t \in [0, 1]$ | gradient coordinate corresponding to the spatial trajectory |
| $\mathcal{G} := (\mathcal{V}, \mathcal{E})$ | heterogeneous spatial graph |
| $\mathcal{V} := \mathcal{V}_x \cup \mathcal{V}_u$ | heterogeneous graph nodes |
| $\mathcal{E} := \mathcal{E}_{xx} \cup \mathcal{E}_{uu} \cup \mathcal{E}_{ux}$ | heterogeneous graph edges within and across modalities |
| $d_{ij} \in \mathbb{R}_+$ | spatial distance for edge $(i, j) \in \mathcal{E}_{xx}$ |
| $\Theta = \{\lambda, \vartheta, \theta\}$ | generative model parameters |
| $\Phi = \{\phi, \psi, \xi\}$ | variational (recognition) model parameters |

#### Multi-modal graph construction

LYNX employs a joint spatial graph structure that facilitates message passing within and across modalities. Given a pair of registered observations from cells 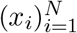 and patches 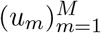, we define a heterogeneous *G* := (*V, E*) with the following default elements: primary (V_*x*_) and auxiliary (*V*_*u*_) nodes, within-modal edges *E*_*xx*_, *E*_*uu*_ and cross-modal edges *E*_*ux*_ (**Fig. 1, Fig. S1a**). Each node is associated with a unique feature vector and spatial coordinate. The granularity of graph node components depends on the data resolution. For example, each primary node *v*_*i*_ ∈ *V*_*x*_ from Xenium corresponds to a single-cell expression. Analogously, with a multicellular auxiliary, each *v*_*m*_ ∈ *V*_*u*_ is represented by a single pixel or spot, while in high-resolution cases such as histology, it may instead correspond to a patch expression summary surrounding a cell.

Then we construct the edge components starting from within-modal settings. For observations with grid-like structures, an unweighted k-NN graph connects each patch node with its direct 1-hop neighbors (e.g. *K* = 8 for a standard cartesian grid). For single-cell modalities, a generic graph connects spatially adjacent nodes within a radius threshold *r*: e.g. *E*_*xx*_ = {(*v*_*i*_, *v*_*j*_) | *d*_*ij*_ ≤ *r*}. Similarly, cross-modal edges are added if an auxiliary node, upon projecting its coordinate to the paired primary domain, is at most *r* distance away from a nearby primary node: 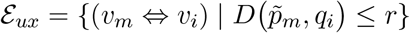, where *p*(·), *q*(·) are coordinate notations to be defined below (see **Methods: Image Registration**). Particularly, the bi-directional edges ⇔ allow distinct edge weights defined in cross-modal pooling / unpooling operators along the LYNX variational / generative stages respectively (**Fig. S1c,e**). In practice, to account for possible registration uncertainties, LYNX sets the radius parameter roughly twice as large as the average mapping ratio across modalities, i.e. on average how many cells are registered to the same patch. It is also possible to choose a cross-modal radius that differs from the within-modal radius, i.e. a larger radius for cell-cell graph encourages longer-range interaction (see **Methods: Hyperparameters**). Because cross-modality coupling is mediated through these local neighborhoods rather than one-to-one correspondences, LYNX is robust to small spatial misalignments between modalities. LYNX implements the hierarchical graph using `pytorch_geometric.data.HeteroData` [84], which enables straightforward subgraph partitioning for batched training.

#### Generative process

We assume paired spatial-omics data acquired from the same specimen or from adjacent tissue sections that have been affinely registered into a common coordinate system (see **Methods: Image registration**) yielding 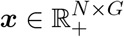 and 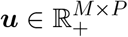.

LYNX defines a conditional generative model

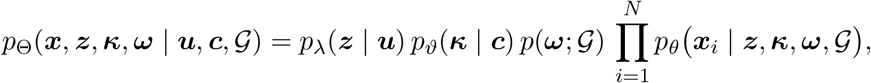

where Θ := {*λ*, ϑ, *θ*} collects all generative model parameters, 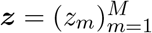 denotes low-resolution niche embeddings, 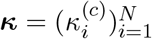 are cell-type specific offsets, and 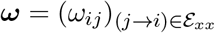 are sparse, distance-aware cell–cell interaction weights defined on within-modality edges *E*_*xx*_ with pairwise distances 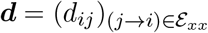. *z, κ*, and *ω* factorize over patches, clusters, and edges, respectively. For notational simplicity, we suppress explicit conditioning on *G* in the following and only write the distances *d* explicitly when discussing the distance-aware priors on *ω*.

##### Auxiliary-anchored niche latents

For each auxiliary spatial unit *m* ∈ {1, …, *M*} (e.g. a DESI pixel or super-cell) we define a conditional prior over its latent niche embedding *z*_*m*_ ∈ ℝ^*D*^:

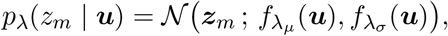

so that 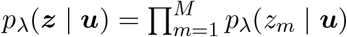. The mean and (diagonal) variance (*µ*_*z*_, *σ*_*z*_) are produced by prior learned neural networks 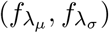. In practice, to construct conditional priors with spatial-awareness, *f*_*λ*_ is parametrized as a single-layer graph convolution networks (GCN) with one-hop message passing between *u*_*m*_ and its direct adjacent patches (**Fig. S1a,d**).

##### Deterministic unpooling to cell-level latents

Given the auxiliary-level latents *z* ∈ ℝ^*M*×*D*^, we obtain initial cell-level latents *s* ∈ ℝ^*N*×*D*^ by a deterministic unpooling operator along the cross-modal edges *E*_*ux*_ (**Fig. S1e**):

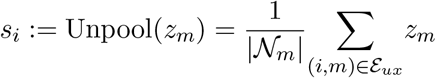

Intuitively, *s*_*i*_ is a łniche embedding” for cell *i* that is anchored in the coarser auxiliary modality while respecting the local cell type composition.

##### Cell-type baseline niche offsets

To represent the basal state of each cell that is interpreted as the absence of neighbor influence, we introduce a cell-type specific latent offset *κ*_*i*_ ∈ ℝ^*D*^:

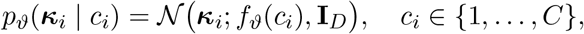

so that 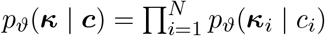. Essentially, each cell *i* is sampled from a Gaussian prior, enabling cell-level variability around its cluster-specific mean *f*_ϑ_(*c*). This can be thought of as learning intrinsic cell-state variability conditioned on the cluster. Here *f*_ϑ_ : {1, …, *C*} → ℝ^*D*^ is a global learnable `torch.nn.Embedding` layer to specify different baseline means per cluster.

##### Distance-aware exponential edge weight priors

For within-modality primary–primary edges *j* → *i* ∈ *E*_*xx*_ with spatial distance *d*_*ij*_ ≥ 0, we define a distance-aware stochastic weight *p*(*ω*; *G*) that encodes the strength of spatial interaction prior from cell *j* to *i*. In the idealized model, we first draw unnormalized edge weights

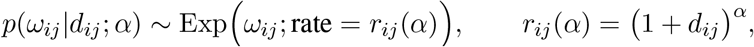

with concentration parameter *α* ≥ 0, and then normalize across all incoming neighbors of *i*:

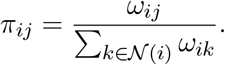

Theorem 1 shows that, when baseline weights do not favor more distant neighbors, the entropy *H*(*α*; *w*) of the normalized weights *π*(*α*; *w*) is nonincreasing with *α* and, in the unique-minimizer case, converges to 0. Corollary 1.1 further implies that the effective number of neighbors *k*_eff_ (*α*; *w*) = exp(*H*(*α*; *w*)) satisfies *k*_eff_ (*α*; *w*) → 1 at an exponential rate as *α* → ∞, i.e. the normalized weights become increasingly sparse and concentrate on the closest neighbor(s). Thus, *π*_*ij*_ and its relaxed counterpart *ω*_*ij*_ quantify a sparse *spatial interaction profile* for each target cell, which can reflect both direct cell–cell signaling and local tissue composition.

##### Distance-aware niche decoupling to single cells

Starting from the deterministically unpooled niche embedding *s*_*i*_ and the cell-type baseline offset *κ*_*i*_, we define the neighbor-transmitted quantity as the baseline-centered residual

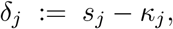

which is mean-centered to *δ*^′^ before message passing to remove shared residual offsets and emphasize local deviations from the global residual baseline. For each receiver cell *i*, we aggregate residual messages from its within-modality neighborhood using the distance-aware edge weights on *E*_*xx*_:

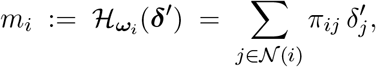

where 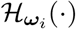 denotes the distance-aware graph filter for receiver cell *i*, and *π*_*ij*_ is defined above as the normalized edge weight. We then form the interaction-updated cell latent as a linear shift to the basal state,

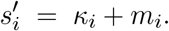

Here 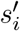 encodes the spatial interaction shift induced by the local neighborhood through 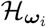 acting on mean-centered niche residuals 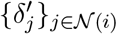 from all 1-hop neighboring cells *j* to the target cell *i*. Formally speaking, the interaction message can be understood as a distance-aware low-pass graph filter applied to the baseline-centered residual field:

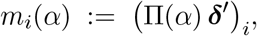

where Π(*α*) denotes the row-normalized, distance-weighted neighborhood operator induced by *E*_*xx*_, i.e., the matrix representation of the graph filter *H*. The full interaction-updated cell embedding then decomposes as

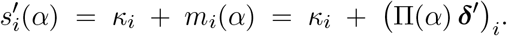

Therefore *H* acts as a row-stochastic low-pass smoother on the mean-centered residual field, and the parameter *α* controls the effective smoothing scale through the distance-decay of *π*_*ij*_(*α*). Empirically, we found that setting *α* = 0.5 as a prior resulted in a balance between meaningful and sparse variational approximation of the *ω* parameter across multiple applications. **Fig. S1e** illustrates the full representation scheme from auxiliary-/niche-level latents through unpooling and distance-filtered message passing to cell-level embeddings.

##### Negative binomial observation model

Given the updated cell embedding 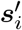, we decode a gene-level mean vector through a feed-forward network *f*_*θ*_ followed by a softmax onto the simplex:

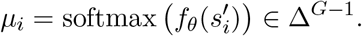

Let *l*_*i*_ denote the library size (total UMI count) of cell *i* and define the expected counts

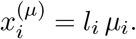

We place a gene-specific inverse dispersion parameter *ρ*_*g*_ *>* 0 collectively with notation 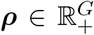. The conditional likelihood factorizes over genes with a negative binomial distribution:

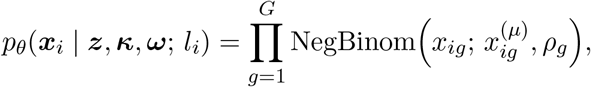

where 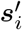 is deterministically computed from *z, κ*, and *ω* through unpooling and the CCI module. The intermediate quantities *s* and *s*^′^ are deterministic functions of *z, κ*, and *ω* through unpooling and the CCI module, so they do not appear explicitly in the likelihood conditioning. The dispersion parameters *ρ* are global parameters shared across cells and learned jointly with the rest of Θ during variational inference.

#### Variational inference

The generative model illustrated above specifies the joint distribution *p*_Θ_(*x, z, κ, ω* | *u, c*) over observations and latent variables we designed: niche embeddings *z*, cell-type baseline offsets *κ*, and cell–cell interaction weights *ω* (**Fig. 1b**; **Fig. S1d**). Computing their exact posteriors is analytically intractable. Therefore, following standard practice in deep generative models [28, 36–38], LYNX introduces amortized approximate posteriors *q*_*ϕ*_(*z* | *x, u*) and *q*_*ψ*_(*κ* | *x*), parameterized by encoder networks with variational parameters Φ := {*ϕ, ψ, ξ*}, while inferring *ω* via a MAP point estimate for computational scalability (see below). All neural network parameters are optimized jointly by maximizing an evidence lower bound (ELBO) on the marginal log-likelihood log *p*_Θ_(*x* | *u, c*) over the primary observation via stochastic gradient ascent and reparameterization trick for *z* and *κ*. The full ELBO decomposes as:

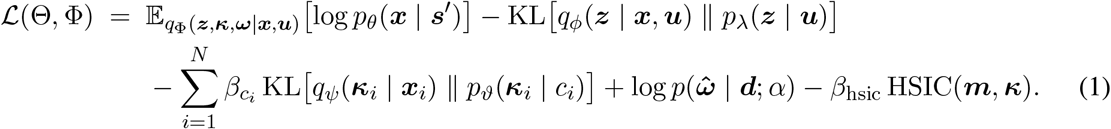

where the expectation is taken over the stochastic variational latents *z* and *κ* and the degenerate MAP posterior over *ω*. The cell embeddings *s* and the interaction-updated *s*^′^ are deterministic functions of *z, κ*, and 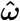 through unpooling and the CCI module, and therefore do not appear as separate latent variables. Similarly, 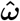 is a MAP point estimate that is deterministic given *z*_cell_ and *x* under *q*_*ξ*_; its contribution enters through the distance-informed prior term log *p*(*ω* | *d*) rather than through a full variational KL term. In addition, the per-cell weight 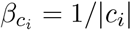 is set as the inverse cell-type frequency with respect to cell *i*, which downweights the KL contribution of *κ* for large cell-type clusters to prevent them from dominating the overall ELBO objective. *m* denotes the residual messages aggregated from the one-hop neighborhood of each cell, and the HSIC term penalizes dependence between *m* and *κ* as described below. All parameters are updated jointly using stochastic gradient optimization.

##### Joint niche posterior q_ϕ_(z | x, u)

The posterior over niche embeddings is parameterized by a cross-modal graph attention network (**Fig. S1b,c**). Primary cell expressions are first projected to *h*_*x*_ ∈ ℝ^*N*×*H*^ via a GCN over *E*_*xx*_, and auxiliary patch features to *h*_*u*_ ∈ ℝ^*M*×*H*^ via a GCN over *E*_*uu*_. A cross-modal `GATConv` then aggregates primary cell embeddings as keys and values into each auxiliary patch node as query over cross-modal edges *E*_*xu*_, yielding a context-enriched patch representation *h*_*m*_ ∈ ℝ^*H*^. Two linear heads map *h*_*m*_ to distribution parameters:

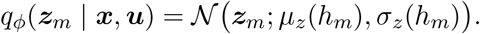

This ensures that each niche embedding conditions on both the auxiliary features of patch *m* and the transcriptomic context of its aligned primary cells, while the posterior remains factorized over patches (**Fig. S1b**). Note that our cell embedding *s* is a deterministic variable without distributional sense, therefore we can directly compute it by applying the previously defined Unpool(·) operator from the posterior *z*_*m*_ here.

##### Cell-type offset posterior q_ψ_(κ | x)

The posterior over cell-type baseline offsets is parameterized by a single-layer MLP applied to log-normalized primary expressions:

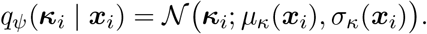

To prevent high-abundance clusters from dominating the corresponding KL term, each cell’s contribution is down-weighted by its inverse cluster frequency 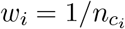, where 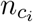 is the count of cells of type *c*_*i*_ in the current subgraph.

#### MAP estimation of interaction weights ω

Rather than placing a full variational posterior over *ω*, LYNX infers each 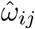 as a MAP point estimate via a 2-layer MLP *g*_*ξ*_:

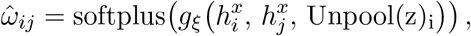

equivalently using a degenerate variational posterior

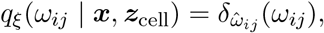

where 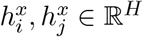 are per-cell embeddings of log-normalized primary expressions through a single-layer MLP, and Unpool(z)_i_ = *s*_*i*_ is the unpooled embedding of the target cell *i*. The MAP design is motivated by both scalability and practical prevention from posterior collapse: a full variational distribution over all | *E*_*xx*_| edges would introduce a huge KL term scaling with edge count | *E*_*xx*_| ≫ *N*, risking overshadowing the reconstruction objective and collapsing *ω* toward the prior. With MAP, the inferred *ω* is driven by the reconstruction signal, while the distance-informed prior serves as a regularizer in the full ELBO (**Fig. S1d**).

##### HSIC independence regularization

To encourage separate representations between the spatial interaction message *m* and the baseline cell-state embedding *κ*, a Hilbert-Schmidt Independence Criterion (HSIC) penalty with an RBF kernel is added to the variational objective. Before constructing *m*, we globally center the transmittable component *δ* = *z*_cell_ − *κ*, so that the resulting message reflects local deviations from the global residual baseline rather than shared residual offsets. Precisely, we define

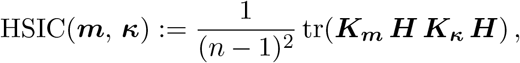

where *n* is the number of cells in the current subgraph partition; *K*_*m*_, *K*_*κ*_ ∈ ℝ^*n*×*n*^ are RBF kernel Gram matrices with entries 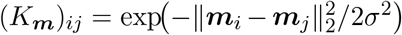 and analogously for *K*_*κ*_; and *H* = *I*_*n*_ − *n*^−1^11^⊤^is the centering matrix [85]. In practice, we compute this empirical HSIC using the algebraically equivalent expanded centered-kernel form, avoiding explicit construction of *H*. We default to a small weight scale *β*_hsic_ = 10^−3^, corresponding to subtracting *β*_hsic_HSIC(*m, κ*) from the ELBO, which intuitively prevents the interaction component from recapitulating cell-type-intrinsic variation already captured by *κ*. In the implementation, gradients through the *κ* argument of the HSIC kernel are detached for stability.

### 2. Cell-Cell Interaction Theory

We provide the following theoretical characterization to identify how the distance-aware prior controls the spatial scale of the inferred cell-cell interactions. Biologically, each receiver cell is assumed to integrate signals from a local neighborhood rather than from the entire tissue. While this assumption does not capture the full context of interactions, we choose to only model these paracrine interactions for simplicity [35]. The normalized weights *π*_*ij*_ can therefore be interpreted as a spatial interaction profile over candidate sender cells, while the entropy of this profile quantifies whether the receiver cell aggregates broadly from many nearby cells or concentrates on a small number of spatially proximal neighbors. The distance-concentration parameters *α* controls this tradeoff: smaller values permit broader neighborhood smoothing, whereas larger values enforce increasingly local communities. This result formalizes the intuition that increasing the distance penalty makes the interaction profile more localized.

#### Theorem 1

(Entropy decay under distance-aware normalization). *Fix a target cell i with finite incoming neighborhood N* (*i*), *and let d*_*ij*_ ≥ 0 *denote the spatial distance from neighbor j to i. Define*

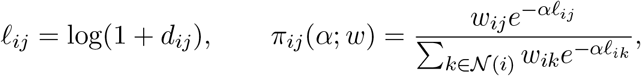

*where w*_*ij*_ *>* 0 *are fixed baseline edge weights and α* ≥ 0 *is the distance-concentration parameter. Let*

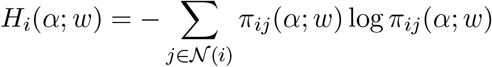

*be the entropy of the normalized incoming edge-weight distribution*.

*Then*

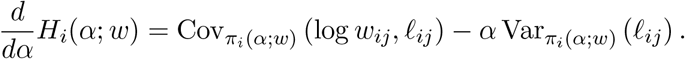

*In particular, if the baseline weights do not favor more distant neighbors, for example if w*_*ij*_ *is nonincreasing as a function of d*_*ij*_, *then*

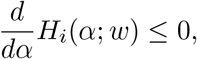

*so H*_*i*_(*α*; *w*) *is nonincreasing in α*.

*Moreover, let*

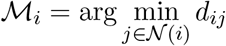

*be the set of closest incoming neighbors. As α* → ∞,

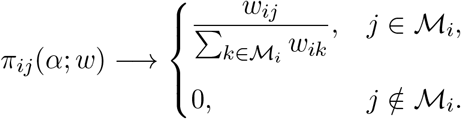

*Consequently*,

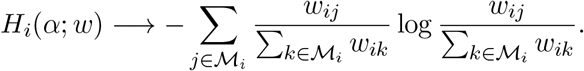

*If the closest neighbor is unique, then H*_*i*_(*α*; *w*) → 0.

In biological terms, Theorem 1 shows that the distance-aware normalization does not merely rescale edge weights; it changes the effective neighborhood size in a predictable direction. When *α* increases, the incoming interaction distribution becomes less diffuse, so the model favors spatially local explanations of cell-state deviations. This is useful for distinguishing broad tissue-composition effects from more localized interaction patterns, because high-entropy profiles correspond to diffuse niche influence, whereas low-entropy profiles correspond to sparse, spatially concentrated interactions.

*Proof*. Write

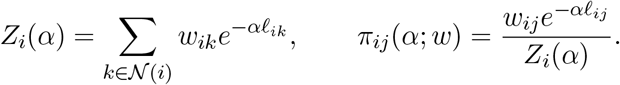

Then

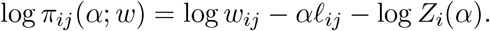

Differentiating gives

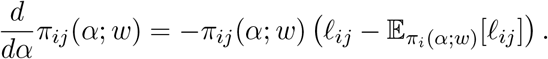

Using

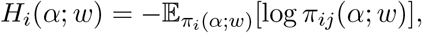

we obtain

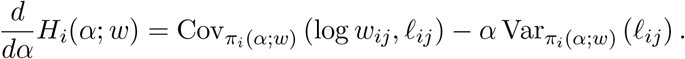

If *w*_*ij*_ is nonincreasing in *d*_*ij*_, then log *w*_*ij*_ is nonincreasing in *ℓ*_*ij*_, so the covariance term is nonpositive, while the variance term is nonnegative. Hence *dH*_*i*_*/dα* ≤ 0.

For the limiting statement, factor out the smallest distance term. Let 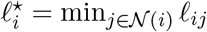. Then

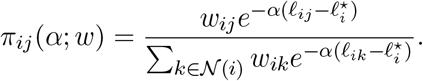

For *j* ∉ *M*_*i*_, one has 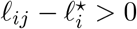, so the corresponding exponential term vanishes as *α* → ∞. Terms with *j* ∈ *M*_*i*_ remain unchanged. This gives the stated limiting distribution and entropy limit.

While entropy provides a mathematical measure of sparsity, it is useful to translate this quantity into an interpretable neighborhood size. We therefore define *k*_eff,*i*_ = exp{*H*_*i*_} as the effective number of incoming neighbors contributing to receiver cell *i*. This quantity equals one when the interaction profile is concentrated on a single neighbor and increases as the receiver aggregates more evenly across multiple neighbors.

#### Corollary 1.1

(Exponential collapse of the effective neighborhood size). *Under the assumptions of Theorem 1, suppose that the closest incoming neighbor of cell i is unique, denoted j*^⋆^, *and define the distance gap*

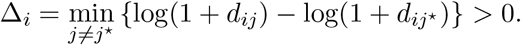

*Let*

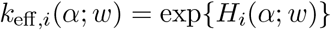

*denote the entropy effective number of incoming neighbors. Then*

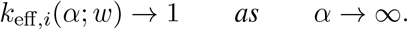

*More precisely, if*

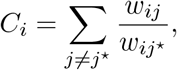

*then the total mass assigned to non-closest neighbors satisfies*

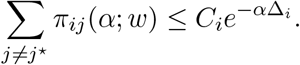

*Consequently*,

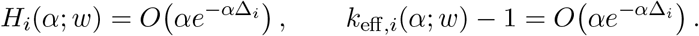

*Equivalently, for any* 0 *< δ <* Δ_*i*_,

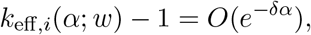

*so the effective number of neighbors collapses to one at an exponential rate*.

*Proof*. Let *j*^⋆^ be the unique closest neighbor and write

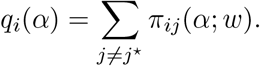

Using the representation from Theorem 1,

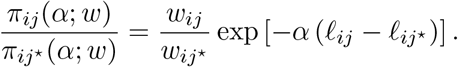

Since 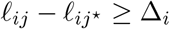 for all *j*≠ *j*^⋆^,

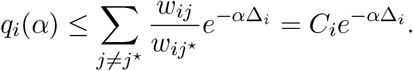

The entropy of a distribution with mass 1 − *q*_*i*_(*α*) on one atom and remaining mass *q*_*i*_(*α*) on all other atoms is bounded by

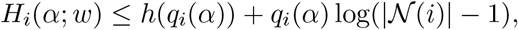

where *h*(*q*) = −*q* log *q* − (1 − *q*) log(1 − *q*) is the binary entropy. Since 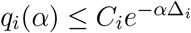, this implies

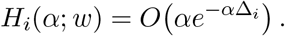

Finally,

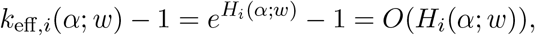

because *H*_*i*_(*α*; *w*) → 0. Therefore

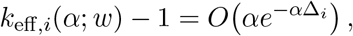

and hence it decays exponentially in *α*.

This corollary gives an interpretable consequence of the distance prior: increasing *α* collapses the effective interaction neighborhood towards the nearest cells. In practice, this supports using *α* as a scale parameter for spatial communication. Smaller *α* values allow LYNX to capture broader microenvironmental context, such as local tissue composition or paracrine fields, whereas larger *α* values emphasize more contact-proximal or highly localized interaction patterns.

Together, Theorem 1 and Corollary 1.1 provide a theoretical basis for interpreting distance-aware normalized weights as a spatial interaction prior. The inferred weights are considered latent interaction strengths regularized by spatial proximity, not direct evidence of a specific ligand-receptor mechanism. We consider this a spatial cell-cell interaction because *ω* deconvolves each local niche into directional contributions from neighboring cells, separating intrinsic cell-type identity from neighborhood-associated residual effects. As we infer interactions on a continuous scale across space, *ω* quantifies how these spatial interaction profiles reorganize across tissue architecture.

### 3 Downstream analysis

#### Spatial gradient inference

After running LYNX, each auxiliary and primary observation units are represented by a smooth latent embedding *z*_*m*_ ∈ ℝ^*D*^, *s*_*i*_ ∈ ℝ^*D*^ respectively. A core interpretable downstream feature of LYNX is its ability to predict spatially-resolved gradients in an unsupervised fashion. To define spatial gradients, LYNX employs principal graph to fit a smooth trajectory *γ* : [0, 1] → ℝ^*D*^ that best summarizes the niche space with a one-dimensional gradient coordinate index *t* ∈ [0, 1]. Conceptually, it constructs a simple undirected graph *G*_*p*_ := (*V*_*p*_, *E*_*p*_) that passes through the latent manifold *z*, minimizing a regularized mean-squared error (MSE) between each latent point *z*_*m*_ and projected principal node *γ*(*t*_*v*_) [86, 87]:

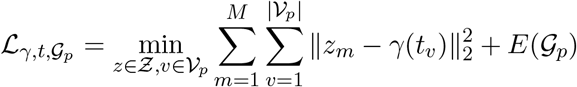

where |*V*_*p*_| specifies the total number of principal nodes, with higher number emphasizing granular manifold structure. *γ* denotes an injection mapping principal graph coordinates onto the embedding space. *E*(*G*_*p*_) regularizes smoothness of the fitted graph detailed below.

##### Principal curve for simple manifold

For tissue samples exhibiting notable, continuous lobular structures, the niche embeddings typically reside on a manifold that can be described by a smooth principal curve without branching points [86]. We reconstruct the graph with `ElPiGraph` [44], which applies an energy-based elastic regularization *E*(*G*_*P*_) := *E*(*µ*_epg_, *λ*_epg_, *R*_0_) to balance curve smoothness with total length. This approach yields a continuous *γ*, extending from “valley” to “peak” endpoints, capturing the dominant biological gradient across the spatial domain. Hyperparameters {*µ*_epg_, *λ*_epg_} control the stretching and bending energy of the elastic curve, whereas *R*_0_ defines the trimming radius when computing the curve-fitting MSE term for each graph node. The principal curve computation is performed by calling `lynx.tl.get_curve`.

##### Principal tree for complex manifolds

For embeddings with more complex geometry such as heterogeneous tumor, LYNX extends the above procedure by learning a principal tree with tips (endpoints) and forks (branching nodes) implemented under `lynx.tl.get_tree`. Precisely, the tree *T* := {*γ*_1_, …, *γ*_*n*_} is estimated using simplePPT implemented with `scFates` [43, 45], where a minimum spanning tree (MST) is refined with graph regularization terms *E*(*G*_*p*_) := *E*(*λ*_ppt_, *σ*_ppt_), with *λ*_ppt_ penalizes the *L*_2_ distance among the fitted graph nodes, and *σ*_ppt_ modulating a negative entropy term measuring the locality of tree nodes. Recommended by [45], we perform a grid search over *σ*_ppt_ to control the tree complexity and with an elbow method to determine the optimal coefficient. LYNX also provides an optional function `lynx.tl.prune_tree(adata, tips_to_keep)` for users to select a subset of tree tips as the desired łroot” and łleaves” to further simplify its topology aligning with biologically interpretable states. The pruned principal tree is then equivalent to *k* principal curves sharing a common root.

##### Projection and gradient coordinate

Given either a principal curve or principal tree, LYNX assigns each niche embedding *z*_*m*_ a scalar coordinate *t*_*m*_ ∈ [0, 1] via orthogonal projection onto the closest point *γ*_*m*_(*t*) along the trajectory. The same procedure is applied to the cell-level embeddings *s* by mapping each *s*_*i*_ to *γ* to obtain its coordinate index *t*_*i*_. An optional annotation (e.g. marker gene) could be specified to rotate the coordinate orientation, where the unsupervised endpoint at *t* = 0 is mapped closer to the marker enrichment. Note that this step only flips the gradient coordinate (*t* → 1 − *t*) without modifying relative patch / cell ordering along the trajectory. For a principal tree, multiple trajectories (*γ*_1_, …, *γ*_*n*_) share the same coordinate system, with *t* = 0 at root note and *t* = 1 at the farthest leaf (terminal) node. The final gradient coordinate iscomputed with `lynx.tl.compute_pseudotime`.

#### Inference of feature dynamics

After integrating the paired observations with a joint spatial gradient, we can interpret it as a spatially-oriented axis by ordering cells and patches along the gradient. For instance, provided a gene expression vector 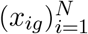, we define its feature dynamic *x*_*g*_(*t*) by ordering the cell indices according to their associated *t*_*i*_ values (**Fig. 1d**). A smoother dynamic can be produced by partitioning *t* into *n* equal-width bins and assigning the average feature expression within each bin. Similarly, provided with cell-type annotations, we compute the phenotype dynamics as the sorted cell-type proportions along the smoothed gradient bins (**Fig. 2h**). A 1D spline regression is used to refit the dynamics for plotting and statistical tests.

#### Discrete zone clustering and abundance test

LYNX further supports clustering the continuous spatial gradient into discrete zones with the Jenks optimization method (jenkspy) [88, 89]. Essentially, taken the 1D gradient coordinate *t* as input, *k* zones with *k* − 1 breakpoints at *t*_0_, …, *t*_*k*−1_ are predicted to minimize the within-zone variance and maximize the cross-zone variance. Given the corresponding auxiliary or primary modality, zone-specific abundance feature modules are further identified with the scanpy function `sc.tl.rank_genes_groups` [90]: a one-sided *t*-test is used for gaussian-like modalities (e.g. DESI) and its non-parametric version Mann-Whitney U test is used for modalities with skewed distributions (e.g. transcriptomics and proteomics). We provide a unified function `lynx.tl.get_zonation_features` accomplishing these steps.

#### Interpretation of cell-cell interaction

The learned per-edge interaction weights *ω* are translated into cell-type-level interaction summaries following model inference. For each target cell *i*, raw weights from its spatial neighbors are first normalized:

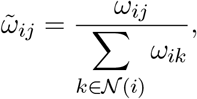

and then averaged by sender cell type *c* to produce a per-cell interaction matrix Ω ∈ ℝ^*N*×*C*^:

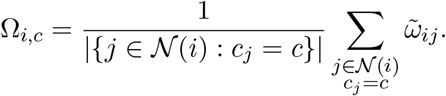

To assess whether inferred interactions exceed what proximity and local cell-type colocalization abundance brings, a distance-weighted null baseline A ∈ ℝ^*N*×*C*^ is constructed. Particularly, for each edge *j* → *i*, the null weight is assigned as 1/(1 + *d*_*ij*_)^*α*^, the expectation value of the exponential prior *p*(*ω*; G); these are normalized per target cell and aggregated by sender type analogously to Ω_*i,c*_. Therefore the baseline *A*_*i,c*_ represents the expected interaction level from local cell-type composition alone, absent any learned co-expression signal.

For each spatial zone and each sender cell type *c*, a one-sided paired-sample *t*-test compares Ω_:,*c*_ against A_:,*c*_ (alternative hypothesis *H*_1_: learned interaction *>* abundance baseline over the gradient coordinate within the defined zone). The resulting − log_10_(*p*) values are reported as node sizes in the zone-specific interaction graphs (**Fig. S3b**, bottom), with edge widths encoding interaction strength and colors indicating the sender cell type (**Fig. 2j**). Zone assignments follow the discrete gradient clustering described above (**Methods: Discrete zone clustering and abundance test**).

### 4 Data preprocessing & analysis

#### Application to human liver data

We apply LYNX to a human liver sample collected in house from a healthy donor. Paired spatial transcriptomics (Xenium), metabolomics (DESI) and antibody staining were collected in consecutive tissue slices 10 µm apart from the left liver region. Human tissue blocks were cryosectioned at 7-µm thickness and placed on Xenium slides (P/N 3000941, Xenium V1, 10x Genomics) for Xenium or charged microscope slide for DESI MSI. For Xenium, sample processing and data acquisition was performed Genome Engineering Core at Columbia University, following manufacturer’s guidelines for fresh frozen samples. Predesigned human Multi-Tissue and Cancer panel with 377 genes (PN 1000626, 10x Genomics) was used. We adopted the Proseg pipeline [91] to refine segmentation and transcript assignment, reporting a total of 67,626 cells. Cell-type annotations were performed with canonical markers and leiden clustering collectively [90]. The antibody staining contains 4 immunofluorescence (IF) zonation-specific markers: GS, CYP, Ass1 and Collagen (**Fig. 2d**). We chose the Xenium sample as the target, and registered both the DESI and IF images against its associated DAPI channel.

DESI MSI was performed in Synapt XS mass spectrometer coupled to Acquity UPLC (Waters Corp) [46]. The tissue was sprayed with solvent with 95:5 methanol: water (v/v) with 0.01% formic acid and 40*pg/µl* leucine enkephalin for positive ion mode and 95:5 methanol: water (v/v) with 0.01% ammonium hydroxide and 40*pg/µl* leucine enkephalin for negative ion mode. Data was acquired at 40 µm spatial resolution. The output data was converted to `imzml` format in HDI software (Waters Corp.) and processed in SCiLS lab software (Bruker, version 2024a) to generate OME-tiff files for data analysis. Data was median normalized and annotated by matching the m/z (mass-to-charge ratio) values with lipidmaps and metabolomics workbench database, resulting in 615 m/z channels in total. The m/z values are matched with the LC-MS data acquired in HDMS mode from the same instrument. Post xenium and DESI analysis, slides were washed with PBS and stained first with anti-CYP3A4 antibody (SantaCruz, Cat# sc-53850) followed by anti-mouse AF750 secondary antibody staining. This was followed by staining the tissue sections with antibodies pre-conjugated with fluorophore: anti-GS-AF647 (Abcam, Cat# ab300751), ASS1-PE (Abcam, Cat# ab210451) and Col1A-AF488 (Cell Signaling, Cat# 28368) and imaging the tissue in PhenoImager HT (Akoya Biosciences).

Quality control filtering excluded Xenium cells with fewer than 20 total UMI counts and genes expressed in fewer than 5 cells. In parallel, image-processing morphological operations discarded overly bright DESI “hotspots” surrounding the tissue edge. Subsequent registration identified the overlapping region between Xenium and DESI measurements, yielding 60,562 cells and 9,564 pixels (patches) that can be mapped to each other across modalities. We set the high-resolution Xenium as the primary modality (*x*) to reconstruct, and the DESI data as the auxiliary *u*. The IF antibody staining is used for held-out validation (see **Methods: Evaluation**).

We performed iterative cell-type annotations for the 60,562 Xenium cells passing quality control and spatial registration against the DESI image. Specifically, we first assigned major cell types based on UMAP visualizations of the log-normalized marker expressions of non-overlapping canonical markers (e.g. `fibroblasts: [‘FBN1’, ‘THY1’, ‘ASPN’]; hepatocytes: [‘CYP3A4’, ‘CYP2A7’, ‘CYP2B6’, ‘APOA5’]`, etc.. Then, we further explored possibilities of fine-tuning cell-states with leiden subclustering on subset cells of each major cell-type. Specifically, we applied `scanpy.tl.rank_genes_groups` function and dotplots to visualize upregulated DEGs defined as adjusted p-value *<* 0.05 (wilcoxon rank-sum test) and logFC *>* 2.0. Given rather limited gene panels (< 400 total genes), we took a relatively conservative approach to only assign refined cell-states with separable marker signals from the dotplots upon subclustering. For instance, we didn’t proceed to distinct “CD4+” and “CD8+” cells from lymphocytes due to the lack of separable signals from the markers; similarly, the cutoff point for “peri-central hepatocytes” and “peri-portal hepatocytes” were unclear as well between *CYP3A4* (peri-central marker) and *CYP2A7* (peri-portal marker), so we kept the broad “hepatocytes” annotation to avoid uncertainty propagation. A full reproducible annotation pipeline is available in the LYNX [notebook example](https://github.com/azizilab/Lynx/blob/main/liver/annotation.py).

For model training, we build the heterogeneous graph with a *k*-NN graph (k=8) for metabolite edges given the cartesian grid nature from regular image patches, and radius-based thresholds for cell-cell edges and cross-modal edges. Raw transcript expressions are used to fit the LYNX decoder reconstruction, but log-normalized RNA counts are fed into the encoder, which is widely applied in attention applications for numerical stability [35, 92]; DESI pixel intensities are rescaled to [0, 1] per channel. For downstream analysis, we empirically categorized the continuous spatial gradient into 4 zones (zone 0 to zone 3): zone 0 denotes the portal triad-proximal region, and zones 1–3 align with the three canonical hepatic metabolic zones [8], corresponding to periportal, mid-lobular, and pericentral territories, respectively. Scanpy function `sc.rank_genes_groups` is used for zone-specific marker selection, with a Mann-Whitney U test used for zero-inflated Xenium, and *t*-test used for gaussian-distributed DESI features (**Fig. S3**).

#### Application to human breast cancer data

We apply LYNX to a human breast cancer sample profiled with Xenium (v1) and post-Xenium H&E on the same formalin-fixed, paraffin-embedded (FFPE) tissue blocks [19]. The tissue (replicate #1) was pathologically reported with HER2+/ER+/PR-receptor status and annotated as invasive ductal carcinoma with a mixture of DCIS and invasive tumor regions. To better understand the molecular distinctions across tumor subtypes, we selected a particular region investigated by Chitra et al., [50], which contains a rich cell type composition including stromal, DCIS, invasive tumor and atypical ductal hyperplasia (ADH). The subset region contains a total of 16,621 cells and 313 genes paired with the histology approximately 425µm × 370µm (2000 × 1750 pixels).

We define the Xenium as the primary modality 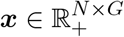 and preprocess the auxiliary histology into patches associated around each cell. First, the original ROI H&E image was z-scaled per RGB channels. Then we extracted *N* = 16, 621 histology patches *u*_*m*_ ∈ ℝ^3×*F* ×*F*^ centered around each Xenium cell, with the patch width *F* = 32 by default. In particular, this allows each patch to roughly contain the histological identity of its own cell and direct neighborhood without over-dilation, i.e. each pixel is on average included in (*N* × *F* ^2^)/(*H* × *W*) ≈ 4.86 patches. The histology patches were flattened to be saved into an expression matrix *u* ∈ ℝ^*N*×(3×*F* ×*F*)^, but would be reshaped back for model training.

Similar to the liver applications, radius-based cell-cell edges *E*_*xx*_ and histology-cell edges *E*_*ux*_ are added to the integrated heterogeneous graph. Nevertheless, as each patch contains overlapped neighborhood information, it is unnecessary to further define *E*_*uu*_. Precisely, in place of graph convolution *across* patches, a stack of two Conv2D layers extract semantic representations *within* each 3 × *F* × *F* patch *u*_*i*_ for both auxiliary embedding paths included in *p*(*z* | *u*) and *q*(*z* | *x, u*) (**Fig. S1a-c**, see **Methods: The LYNX model**). Broadly speaking, this minor architecture modifications could be generalized in integrations where the auxiliary contains high-resolution grid structure with extremely shallow feature throughput (e.g. histology image with RGB-channels).

#### Application to mouse thymus data

We apply LYNX to a spatial multi-omics data from a mouse thymus section collected by Liao et al. [71]. Stereo-CITE-seq jointly profiles whole-transcriptome mRNA and a 51-plex panel of cell-surface proteins from the same tissue section under bins with 50µm × 50µm resolution without need for registration. Naturally, we assign the RNA expressions as the primary modality given its much richer feature dimensions.

For the mRNA expression, we applied scanpy preprocessing functions to select the top 3,000 highly-variable genes plus canonical cell-type markers after removing ribosomal genes. Additionally, low-quality bins with fewer than 10% expressed genes were removed, which we found spatially corresponding to the technical artifact regions outside the tissue capsule [26]. For the protein expressions, we normalized their raw intensities to a feasible range (*u* ∈ [0, 24, 103] → *u*^′^ ∈ [0, 8.50]) with an inverse hyperbolic function [20, 93]: *u*^′^ = arcsinh(*u/c*), where the cofactor *c* defaults to 10. The paired RNA and protein expressions shared the same bin coordinates without registration needs. After data preprocessing, we saved them under two `AnnData` objects with a total of 4,169 bins.

Given the cartesian grid nature of the multicellular bins, a simple k-NN spatial graph *k* = 8 is constructed for both within- / cross-modal edges. Similar to the human liver application, log-normalized RNA expressions are passed to the LYNX encoder. In addition, since the bin resolution doesn’t contain single-cell information, LYNX bypasses the sparse message passing decoupling steps, and directly reconstructs the raw RNA expressions *x* from the cell-type unaware latent *s*. For model validation, we annotated the capsule and medulla layers with marker-based hand-gating, and further constructed a ground-truth cortico-medullary axis (CMA) and spatial clusters following OrganAxis [26], a marker-based annotation tool that reconstructs the spatial gradients of capsule → cortex → cortico-medullary junction (CMJ) → medulla with a nonlinear distance transformation (**Fig. 4a**, see **Methods: Evaluation**).

#### Image registration

Upon data collection, spatial-omics from adjacent sections are often subject to tissue placement distortions, structural changes and sample preparation artifacts (e.g. tearing, folding, etc.). A few end-to-end whole-slide alignment approaches attempted to address these challenges [94–96]. Nevertheless, trivial registration automation across different imaging platforms is practically infeasible due to inconsistent landmark detections. To accurately map expressions from different modalities onto a common coordinate system, we propose an interactive pipeline with simple affine transformation tailored by recent data structure advancements such as `anndata` [97] and `napari-spatialdata` [76, 77].

##### 2D registration

For a given pair of slices, we define one modality as the source (moving) and the other as the target (fixed) along with their associated images. Each observation is first converted to a `SpatialData(sdata)` object, where the molecular expressions, their corresponding coordinates and the full image are saved. First, the source and target objects are jointly loaded into a napari viewer, where users can interactively annotate sets of moving and fixed landmarks {(*a*^*s*^, *b*^*s*^)}, {(*a*^*t*^, *b*^*t*^)} from conserved anatomical structures. We recommend marking at least 5 paired anchors for robust registration across platforms and resolutions. With these landmarks, we then estimate a 2D affine transformation *M* ∈ ℝ^3×3^, and compute the transformed source coordinates with 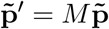, where each original source coordinate 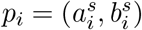 is augmented as a homogeneous column vector 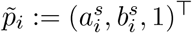. In parallel, the target coordinates 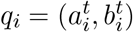 could be inversely mapped to the source domain by applying 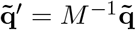.

Finally, we overwrite the warped source image under `sdata[‘image’]` and update the transformed source and target expression coordinates under each `sdata[‘table’].obsm`. Notably, all the affine transformation parameters are saved under `sdata` objects as well, which supports fast querying of either modality in the other’s domain.

##### 3D registration

The same registration pipeline can easily extended to serial sections profiled with alternating platforms (**Fig. 5a**). Here we introduce the approach with a simplified scenario, where the spatial-omic (*x*^(*l*)^) is collected at *L* equally spaced depths (0, …, *l* − Δ*d, l, l* + Δ*d*, …, *L*) and a secondary paired modality (*u*^(*l*+1)^) at each *l* + 1 layer. We adopt a two-step procedure: first, we treat modality 1 as the backbone by selecting a middle layer *l*^∗^ as the target slice and iteratively aligning its neighbors *l*^∗^ ± Δ*d, l*^∗^ ± 2Δ*d*, … to the most recently warped slice, thereby propagating a consistent affine transform along the depth axis:

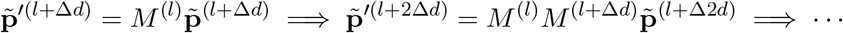

Second, for each depth *l*, the corresponding modality 2 slice at *l* + 1 is registered onto the warped modality 1 slice at *l*, ensuring that both modalities share a common 3D coordinate frame. In practice, the higher-resolution modality is always chosen as the target frame / backbone.

A full pipeline for paired Xenium-DESI registration could be found in the LYNX [notebook example](https://github.com/azizilab/Lynx/blob/main/liver/alignment.py); specific helper functions are accessible from `lynx.tl.utils.registration`.

### 5. Hyperparameters

#### Global settings

Across all biological applications, the LYNX model training utilizes a consistent core architecture. The encoder and decoder networks are configured with a hidden dimension of *H* = 32 and a bottleneck latent dimension *D* = 6 (**Fig. S1b**). SiLU (Sigmoid Linear Unit) is used as the activation function. To facilitate memory-efficient training, spatial heterogeneous graphs were randomly partitioned into 16 subgraphs using `pytorch_geometric.ClusterData` akin to mini-batches in tabular data. All models were optimized using the Adam optimizer with a weight decay of 1 × 10^−3^ and (*β*_1_, *β*_2_) = (0.95, 0.999). Training was conducted for a maximum of 500 epochs with a 7:3 split between training and validation sets.

#### Application-specific hyperparameters

While the core architecture remained stable, hyperparameters governing graph topology and modality-specific optimization were tailored to each tissue type (Table 1). Specifically, the learning rate and patience were adjusted to account for the varying signal-to-noise ratios across spatial metabolomics (DESI), proteomics (CITE-seq), and imaging (H&E) modalities. Empirically, for healthy samples with a simple latent trajectory assumption, we found training converges quickly with a larger learning rate and a progressive early-stopping scheme.

**Table 1:** Summary of application-specific model hyperparameters.

| Application | Graph parameters | Learning rate | Patience | CCI Module |
| --- | --- | --- | --- | --- |
| Human liver | $k = 8, r = 50$ | $1 \times 10^{-2}$ | 20 | ✓ |
| Mouse thymus | $k = 8$ | $1 \times 10^{-2}$ | 20 | ✗ |
| Human breast | $F = 32, r = 50$ | $1 \times 10^{-3}$ | 20 | ✓ |

Next we highlight the guidelines for the downstream principal graph inference on the joint latent manifold (see **Methods: Spatial gradient inference**). A perfect trajectory fitting requires hyperparameter tunings specific to particular embedding manifold, we report the following default value ranges experimentally robust (Table 2). For a simple principal curve fitting with `Elpigraph`, a total of 20 principal nodes is sufficient without overfitting a complex trajectory. The elastic stretching parameter is fixed at *µ*_epg_ = 1.0, with the bending parameter *λ*_epg_ varying between 0.1–0.01 (less bending regularization for a manifold with more data points) is empirically satisfying. For a more complex principal tree fitting with `simplePPT`, the number of nodes defaults to the 1% of the total data points; the optimal bandwidth parameter is obtained with a grid search between 1-0.01, whereas the length penalty term is encouraged to be set larger than 10^2^ to avoid squiggled tree paths (overfitting).

**Table 2:** Hyperparameter description for spatial gradient inference.

| Symbol | Description |
| --- | --- |
| n_nodes | Number of nodes in the principal graph |
| epg_mu ($\mu_{\text{epg}}$) | Elasticity "stretching" parameter for principal curve (↑ for spreadness) |
| epg_lambda ($\lambda_{\text{epg}}$) | Elastic "bending" parameter for principal curve (↑ for less curvature) |
| trim_radius_ratio ($R_0$) | Trimming radius ratio for outlier robustness (↑ for less outlier consideration) |
| ppt_sigma ($\sigma_{\text{ppt}}$) | Bandwidth parameter for principal tree (↑ for complex branches) |
| ppt_lambda ($\lambda_{\text{ppt}}$) | Length penalty term for principal tree (↑ for shorter tree) |

### 6 Evaluation

To evaluate the robustness of spatial gradient predictions, we construct biologically informed ground-truths for the liver and thymus applications, where the known “porto-central-axis” and “cortico-medullary-axis” are well studied in previous literature [2, 26, 79]. Here we outline the important evaluation steps.

#### Ground-truth annotation

##### Thymus: cortico-medullary axis

To generate the ground-truth cortico-medullary axis (CMA) and discrete layers (Capsule, Cortex, CMJ, Medulla), we implemented a spatial reconstruction pipeline based on an established marker-based annotation framework called OrganAxis [26] (**Fig. 4a**). First, we identified representative spatial cores for the capsule, cortex, and medulla through manual gating. *CD4* marks the cortex, and a combination of *CD29* and *Cd45* was used to delineate the capsule, the region that harbors the most concentrated immature thymocytes. While canonical medullary markers (e.g. *CD5* and *CD44*) exhibited poor signals in both proteomics and transcriptomics profiles, the medulla was localized using the negative expression of CD8a.

Next, we employed the OrganAxis approach to compute a continuous spatial coordinate framework defaults to *t* ∈ [−1, 1]. The axis is defined by potential functions *H*(*A, B*) based on the relative proximity to tissue structures *A* and *B*:

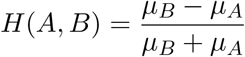

where *µ*_*s*_ denotes the mean Euclidean distance to the *k*-nearest neighbors within structure *S*. The final CMA value was computed as a weighted combination:

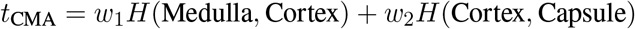

effectively mapping the tissue from the subcapsular zone through the cortex to the medullary core. We followed the default settings (*w*1 = 0.8, *w*2 = 0.2, *k* = 10) reported in the original paper, where Yayon et al. [26] computed for mouse thymus samples from Visium/IBEX platforms. Lastly, the continuous CMA was binned into functional clusters (`Cortico-Medulla Layers`) with empirical thresholds: Capsule (*t*_CMA_ *<* −0.2), Cortex (−0.2 ≤ *t*_CMA_ *<* 0.2), Cortico-Medullary Junction (CMJ, 0.2 ≤ *t*_CMA_ *<* 0.45), and Medulla (*t*_CMA_ ≥ 0.45). We finally normalized the ground-truth CMA scores to [0, 1] for our benchmarks, with capsule closer to 0 and medulla closer to 1.

##### Liver: porto-central axis

Similarly, to establish the ground-truth porto-central axis (PCA) for the liver analysis, we employed a hybrid approach combining empirical histological annotation with a biophysical model of metabolic zonation. First, we identified the spatial boundary conditions by annotating the PV & CV regions based canonical markers from the aligned antibody staining: PV(*Col*), PP(*Ass1*), PC(*CYP*), CV(*GS*) (**Fig. 2d**). Cells identified within these vascular cores were assigned fixed Dirichlet boundary values for the axis reconstruction. Specifically, we set the PV cells as 0, CV cells as 1, and the cells bordering PP & PC as 0.5.

To generate the continuous axis values for the intermediate parenchyma, we adopted the “metabolic uptake” model proposed by Karschau et al [48]. This framework postulates that the zonation profile is governed by the steady-state distribution of metabolic substrates, balanced by tissue transport and cellular consumption. Simplifying the transport dynamics to a diffusive process, the concentration field *u*(x) is modeled by a reaction-diffusion equation:

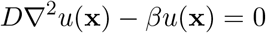

where *D* is the effective diffusion coefficient representing intercellular connectivity, and *β* represents the rate of metabolic uptake. Unlike simple distance-based metrics, this model accounts for the non-linear decay of molecules as they traverse the hepatic lobule with blood or bile flows. We solved this system numerically on the discrete cellular landscape with a spatial connectivity graph constructed using Delaunay triangulation of cell centroids. The continuous partial differential equation was then reformulated as a sparse linear system involving the graph Laplacian *L* [47, 98]:

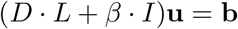

where *I* is the identity matrix and b encodes the aforementioned boundary constraints: b_PV_ = 0, b_CV_ = 1 and b_PP/PC border_ = 0.5. The system was solved for the rest of nodes u_*i*_ ∈ [0, 1] as the continuous ground-truth spatial gradient coordinate *t* along the porto-central axis (**Fig. S2a**).

#### Metrics

Given the ground-truth coordinate indices *t*^gt^ along the spatial axes of liver and thymus, we benchmarked the predicted spatial gradients *t*^pred^ from LYNX against competing methods. A range of regression- / classification- / autocorrelation-based metrics were applied as follows:

##### Spearman’s Correlation Coefficient

We applied Spearman’s correlation to assess the monotonic relationship between the ground-truth gradient coordinates and predictions while account for possible non-linear scalings:

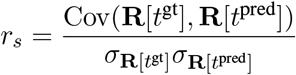

A higher value indicates a better cell ordering preservation **R**[·] along the true spatial gradients.

##### Root Mean-Squared Error (RMSE)

Besides the global correlation coefficient, we quantified the average m_J_agnitude of deviation between the predicted gradient coordinate values and the ground-truth: 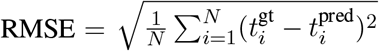. Lower RMSE values indicate a higher accuracy of the inferred trajectory in recovering the exact values of the underlying spatial gradient.

##### Average Precision (AP)

To fully leverage the held-out antibody staining validations, we can reformulate the gradient coordinate values *t*^pred^ as prediction scores classifying cells into spatial zones defined by antibody markers. Precisely, ground-truth binary labels were generated by thresholding antibody expressions of each channel, where 1 represents “central-like” regions. In particular, for markers staining portal regions, we inverted the original binary labels so that a label of 1 always corresponds to łpericentral”. The overall AP score provides a summary of classification power at different levels of complexity: e.g. predicting positive labels for *GS* (exact central vein regions) vs. predicting positive labels for *CYP* (pericentral regions). Complementary to AP, we also plotted the Area Under the Receiver Operating Characteristic Curve (ROC-AUC) to pericentral and periportal regions, respectively (**Fig. S2c**).

##### Clustering Metrics

We extended the continuous gradient benchmark to a spatial clustering task using mouse thymus data, where we evaluated how accurately the capsule, cortex, CMJ, and medulla regions were localized. Followed by a preliminary hungarian matching algorithm finding the optimal mapping between each pair of ground-truth and predicted cluster labels 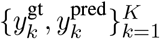, we evaluated the ratio of their cluster assignments with Adjusted Rand-index (ARI) and two adjusted mutual information (NMI, AMI) (**Fig. S7b**).

##### Moran’s I

We further measured the spatial autocorrelation of each spatial gradient prediction to quantify their smoothness. Moran’s I is defined as

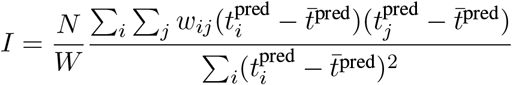

where *N* is the number of cells, 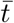 is the mean gradient value, *w*_*ij*_ ∈ {0, 1} is the spatial connectivity between cells *i* and *j*, and *W* is the sum of connectivity degrees. The spatial graph was constructed based on Delaunay triangulation with `squidpy.gr.spatial_autocorr`. Higher Moran’s I indicates better spatial coherence rather than exhibiting random noise along the predicted spatial axis.

#### Ablation studies

To dissect the contribution of the LYNX architecture, we performed a series of ablation studies on the human liver and mouse thymus datasets with ground-truth spatial gradients. Specifically, we wanted to show that the auxiliary modality is actively regulating the predicted latent embeddings and consequently on the spatial gradient inference. First, we built a reduced model without the auxiliary modality (*u*) and multi-modal integration, where the model only reconstructs the primary modality *x* from the latent *z* with a single-omic VAE architecture with graph smoothing encoders. In addition, we constructed a noise-perturbed baseline: LYNX’s conditional generative and multi-modal integration variational paths are kept, but spatial coordinates of *u* are randomized to break the correct local alignment across modalities. The full ablation results are available in **Fig. S8**, which revealed entangled latent representations and performance drops from the reduced/perturbed baselines, highlighting the importance of spatially informative multi-modal integration.

#### Benchmark

We benchmarked LYNX against recent state-of-the-art methods for multi-omics integration and spatial gradient predictions. For methods generating end-to-end spatial gradients, we directly output their 1D gradient coordinates. Alternatively, for methods that produce or integrate latent representations, we further computed the spatial gradient inference applied to LYNX, with hyperparameters fine-tuned individually. For the liver application, we set *k* = 30 for methods requiring spatial k-NN graph construction (SIMVI, spatialMETA),(roughly the average neighbors to our *r* = 50). For the thymus application, we set a global *k* = 8 for the grid-structure Stereo-CITE-seq across all benchmarked methods. We report the inferred spatial gradient coordinate 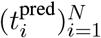 with respect to the primary omic data, rotate the orientation with the same endpoint marker(s) and rescale it to [0, 1] for parallel comparisons. All experiments and runtime measures were conducted on the same machine with 24-core AMD Ryzen9 3900X CPU and a GeForce RTX 2080 GPU. The specific implementation and hyperparameters used in each method are detailed below:

- **GASTON** We directly obtained the GASTON 1D spatial gradient prediction (“isodepth”) following the tutorial https://gaston.readthedocs.io/en/latest/notebooks/tutorials/xenium_brca_tutorial.html. Given the transcriptomics expression matrix, we computed the GLM-PCA values as model input, trained the neural networks with the default 30 random seeds, 2 hidden layers, 20 neurons per hidden layer and 100,000 epochs to ensure its convergence. The best-performing model was used to compute the isodepth with `gaston.dp_related.get_isodepth_labels` before scaling to *t*^GASTON^ ∈ [0, 1].
- **SpaceFlow** We followed the OmicVerse package [99] to compute the 1D pseudo-Spatial Map (pSM): https://starlitnightly.github.io/omicverse/Tutorials-space/t_spaceflow/. We trained the model with `ov.space.pySpaceFlow.train` and default hyperparameters: 50 latent dimensions, 1,000 total epochs. The 1D pseudo-Spatial map (pSM) was then extracted with `ov.space.pySpaceFlow.cal_pSM`.
- **SIMVI** We followed their MERFISH tutorial with default hyperparameters to compute the spatially informed embedding: https://idsc-suite.readthedocs.io/en/latest/simvi/tutorials/SIMVI_tutorial_MERFISH.html. In particular, SIMVI jointly learns an intrinsic embedding `simvi_z` ∈ ℝ^*N*×20^ and a spatial variation embedding `simvi_s` ∈ ℝ^*N*×20^ with simvi.model.SimVI.train. To extract the embedding interpretable for spatial gradients, we saved `simvi_s` for computing the final gradient coordinate values.
- **Novae** We fine-tuned on our spatial human liver and mouse thymus data with the pretrained models `novae-human-0` & novae-mouse-0, respectively: `model=Novae.from_pretrain; model.fine_tune`. We then saved the predicted latent representation defaulting to 64 dimensions for spatial gradient predictions with `model.compute_representation`. The full tutorial is available at https://mics-lab.github.io/novae/tutorials/main_usage/. The zero-shot inference results without fine-tuning were also reported.
- **scVI & TotalVI** We trained the multi-omic integration approach TotalVI with the default `n_latent = 20` and 400 total epochs. Since TotalVI requires paired-omics data with the same resolution (*M* = *N*), to benchmark on the human liver dataset, we upsampled the pixel-level DESI by interpolating the expressions to their closest Xenium cells following image registration. The joint latent embedding was extracted from `scvi.model.TOTALVI.get_latent_representationto` compute the spatial gradients. We also computed the single-omic baseline results with scVI. The full tutorials are available at https://docs.scvi-tools.org/en/1.3.3/tutorials/.
- **SpatialGlue** We computed the spatially-informed joint embedding from their multi-omic integration pipeline https://spatialglue-tutorials.readthedocs.io/en/latest/. In the human liver application, we interpolated the low-resolution DESI spots to the paired high-resolution Xenium cells. Noticeably, training on large-scale Xenium data consistently ran out of memory with the smallest possible k-NN graph setup (*k* = 3), we therefore were only able to include their results for the thymus benchmark.
- **spatialMETA** We followed the official tutorial https://spatialMETA.readthedocs.io/en/latest/tutorials/ to compute the joint spatial embedding. To compensate for resolution differences, we applied the function `spot_align_byknn` to interpolate lower-resolution spatial-modality to map the higher-resolution in a unified coordinate space.

In addition, we also benchmarked against classic dimension reduction and pseudotime inference methods including PCA, Diffmap and DPT, where the implementations could be easily reproduced from scanpy tutorials [90].

## Data Availability

Raw and preprocessed data for the liver studies are accessible via Zenodo: https://doi.org/10.5281/zenodo.21110855. The raw thymus data (sample *Mouse_Thymus1*) from Liao et al. [71] is publicly available at https://zenodo.org/records/10362607. The raw breast cancer is publicly available on the official website of 10X Genomics: https://www.10xgenomics.com/products/xenium-in-situ/ preview-dataset-human-breast. The processed AnnData or SpatialData objects corresponding to each application are also available upon request.

## Code Availability

The LYNX package and sample tutorials are available via GitHub with BSD3 license https://github.com/azizilab/Lynx. Reproducible pipelines for generating the paper figures are available at https://github.com/azizilab/Lynx_reproducibility.

## Acknowledgements

We thank Bianca Dumitrascu, José L. McFaline-Figueroa, Mingxuan Zhang, Aaron Zweig, Justin Hong, Khushi Desai and Nicolas Beltran-Velez for their insightful discussions and feedback. Y.J. acknowledges support from the Columbia University Presidential Fellowship; J.D.M acknowledges support from the Columbia University Blavatnik Fellowship; J.S.M acknowledges from the Columbia University Guthikonda Fellowship. The research of B.R.S. is supported by the National Cancer Institute (P01CA291697), NIH 4UH3CA256962), a Cancer Center Support Grant (P30CA008748), and the Columbia University Digestive and Liver Disease Research Center (funded by NIH grant 5P30DK132710) through use of its Bioimaging Core, Bioinformatic and Single Cell Analysis Core, Organoid and Cell Culture Core and Clinical Biospecimen and Research Core, and the Cancer Center Flow Core Facility and Genomics and High Throughput Screening Shared Resource funded in part through the National Institutes of Health/National Cancer Institute Cancer Center Support Grant (P30CA013696). B.R.S. and E.A. acknowledge support from a Columbia University HICCC-SEAS seed grant. E.A. is supported by the NIH NHGRI grants R01HG012875, R21HG012639, NSF CBET 2144542, grant number 2022-253560 from the Chan Zuckerberg Initiative DAF, an advised fund of Silicon Valley Community Foundation, and the Africk Family Fund.

## Author contributions

Y.J., J.D.M., P.R. B.R.S and E.A. conceived the project. Y.J., J.D.M. and E.A. designed and developed the LYNX model. P.R. conducted the sample collections, spatial-omics experiments and result interpretations. J.Y.Z. and K.F. contributed to model implementation and benchmarking. J.S.M and N.H. contributed to the analysis pipeline of multi-modal spatial alignment. B.R.S. and E.A. supervised the project and provided critical feedback on the model design and biological interpretations. All authors contributed to the writing of the manuscript.

